# Manipulation of natural transformation by AbaR-type islands promotes fixation of antibiotic resistance in *Acinetobacter baumannii* populations

**DOI:** 10.1101/2023.10.06.561211

**Authors:** Rémi Tuffet, Gabriel Carvalho, Anne-Sophie Godeux, Maria-Halima Laaberki, Samuel Venner, Xavier Charpentier

## Abstract

The opportunistic pathogen *Acinetobacter baumannii*, a major global public health concern, has developed multiple variants of AbaR-type genomic islands that confer multidrug resistance. The mechanisms facilitating the persistence of these potentially costly islands within *A. baumannii* populations have remained enigmatic. In this study, we employed a combination of investigative methods to shed light on the factors contributing to their selective advantage and long-term persistence. The dissemination of AbaR islands is intricately linked to their horizontal transfer via natural transformation, a process through which bacteria can import and recombine exogenous DNA, facilitating allelic recombination, genetic acquisition, and deletion. In experimental populations, we first quantified the rate at which natural transformation events occur between individuals. Our findings indicate that the rate of AbaR deletion events is marginally higher than the rate of their acquisition. When this result is integrated into a model of population dynamics not exposed to antibiotic selection pressure, it leads to the swift removal of AbaRs from the population, a pattern that stands in contrast to AbaR prevalence in genomes. Yet, genomic analyses show that nearly all AbaRs-carrying *A. baumannii* have at least one AbaR disrupting *comM*, a gene encoding a helicase critical for natural transformation. We discovered that such disruption differentially inhibits the rate of genetic acquisition and deletion. Specifically, they significantly impede the removal of AbaRs while still enabling the host cell to acquire and recombine short sequences, such as allelic variants. Through mathematical evolutionary modeling, we demonstrate that AbaRs inserted into *comM* gain a selective advantage over AbaRs inserted in sites that do not inhibit or completely inhibit transformation, in line with the genomic observations. The persistence of AbaRs within populations can be ascribed to their targeted integration into a gene, substantially diminishing the likelihood of their removal from the bacterial genome. In contrast, this integration enables the host cell to preserve the ability to acquire and eliminate various short heterologous sequences, enabling the host bacterium - and thus its AbaR - to adapt to the unpredictability of the environment and persist over the long term. This work underscores how AbaRs, and potentially other Mobile Genetic Elements (MGEs), can manipulate natural transformation to ensure their persistence in populations, ultimately leading to the high prevalence of multidrug resistance.

## Introduction

*Acinetobacter baumannii* is responsible for a wide range of nosocomial infections (1). Due to the constant worldwide increase in its multi-drug resistance (MDR), including to antibiotics of last resort (carbapenems, colistin) (2), *A. baumannii* has been declared critical for research efforts by the WHO (3) and is considered as a major threat to human health by the CDC (4). *A. baumannii* is thought to be a rapidly evolving species, showing high rates of recombination and horizontal gene transfer (HGT) (5–7). Indeed, clinically-relevant resistance levels are largely the result of horizontally acquired antibiotic resistance genes (ARGs) which are hitchhiking on mobile genetic elements (MGEs) (8, 9). Widespread carriers of ARGs and transposable elements (TEs) are the transposon-derived and closely-related genomic islands AbaR-type and AbGRI-type (hereafter collectively called *A. baumannii* resistance islands, AbaRs) (10–12). Examination of AbaRs shows that they result from a complex history of multiple successive transposition events taking place within an original Tn7-like backbone (12–14). AbaRs often contain a class 1 integron (14, 15) and additional modules carrying ARGs conferring resistance to aminoglycosides, tetracycline and sulfonamide and beta-lactams (10). The intense transposition activity driving AbaRs plasticity suggests that they represent a safe harbor and reservoir for diverse TEs and ARGs (10). This plasticity could, for example, be behind the recent emergence of the AbaR4 variant, carrying the carbapenem resistance gene *bla*_OXA-23_, which is a major concern for the evolution of antibiotic resistance in *Acinetobacter* pathogenic species (16, 17).

AbaRs can be considered highly successful genomic islands, as they can be found in most of clinical isolates of *A. baumannii*. This could be due to their ability to transfer across bacteria, by hitchhiking on plasmids (10, 11) but also by direct transfer through homologous recombination (18, 19). The genomic signature of these transfer events, and the fact that most *A. baumannii* isolates are competent for natural transformation (20–22) hint at a major role of this process in their HGT. Natural transformation (hereafter called ‘transformation’) is the process by which bacteria actively import exogenous DNA to recombine it in their chromosome (23). Confirming genomic evidence, transformation was found to allow the transfers of AbaRs at high rates between individual cells of mixed populations (24). Although AbaRs could use transformation for their dissemination, the contribution of transformation in the dynamics of MGEs, and more broadly in genome evolution, is currently questioned (25–27). It is well accepted that transformation can serve as a mean for genetic diversification, allowing the acquisition by recombination of genetic polymorphisms (allelic transfer) and the efficient adaptation to fluctuating environments (28–30). Because transformation allows import of large DNA fragments, it also permits acquisition of large non-homologous DNA sequences (gene transfer) if these are flanked by sequences homologous between donor and recipient genomes (31–34). A consequence is that transformation opens the door to the acquisition of MGEs, such as transposon and integrons, prophages and genomic islands (35–37) which often encode adaptive traits (antibiotic resistance). Yet, MGEs are associated with a fitness cost and, when not conferring an adaptive advantage, bacteria could use transformation as a mean to delete these MGEs from the chromosome (27). Alternating the acquisition and removal of MGEs, and thus the transient expression of MGEs-carried traits, would maximize fitness by buffering against unpredictable environmental fluctuations (38).

Although beneficial in the long term for bacteria, alternating MGEs is not compatible with the propagation and persistence of specific MGEs. Thus, a form of genetic conflict may exist between bacteria and MGEs. In that sense, a growing list of MGEs are found to counteract transformation through diverse strategies (39–45), suggesting an evolutionary response of MGEs against the cleansing activity of bacteria from transformation. However, the consequences of transformation inhibition on the evolutionary success of MGEs remains underappreciated. Yet, it would be critical to the persistence and dynamics of ARGs. In this study, we examine the strategy of AbaRs islands of *A. baumannii*. which are commonly discovered inserted within the *comM* gene (11). This gene encodes a helicase that plays a specific role in the recombination of DNA obtained through transformation (46). As a result, AbaRs could impede the process of transformation. We here test the hypothesis that the interference of AbaR with transformation, mediated by their insertion site, is a strategy that gives them a selective advantage. We first confirmed the almost exclusive insertion of AbaR in *comM* sites among a large number of potential sites. We then quantified the cost of this insertion in terms of growth of the bacterial host population. In addition, we experimentally quantified the effects of inserting AbaR into different genes on the efficiency of transformation (acquisition and elimination of heterologous sequences) in realistic populations. This experimental work was used to develop and parameterize an evolutionary model in which AbaRs that differ in their strategies (insertion site) compete in exploiting the bacteria.

## Results

### AbaR prevalence is associated with insertional inactivation of *comM*

An extensive query of AbaRs in publicly available genomes revealed that over 66% of isolates carry such island (11). Mainly present in the chromosome (85%), they were found inserted at 51 different backbone loci and in 12 loci within other MGEs (prophage, IS). Considering that over half of isolates carry more than one AbaR, we found that 96% of isolates carry at least one inserted in *comM* (Figure 1A). So, while AbaR can occupy diverse insertion sites, these essentially occur when one AbaR is readily inserted in *comM* (Figure 1A). Interestingly, insertional inactivation of other genes essential for natural transformation (*pilT, pilS*) also occur but are rare. In the very few situations when AbaR are not inserted in *comM*, they are mainly found in plasmids (Figure 1A). So AbaRs are either chromosomal with at least one insertion in *comM*, or to a lesser extent, plasmid-borne. Overall, AbaRs show an apparent strong affinity for the *comM* gene which appears as a requirement for occupancy of other chromosomal insertion sites. This suggests that inactivation of *comM* is highly beneficial for AbaRs, i.e. for their long-term fixation in the *A. baumannii* genome.

**Figure 1:**
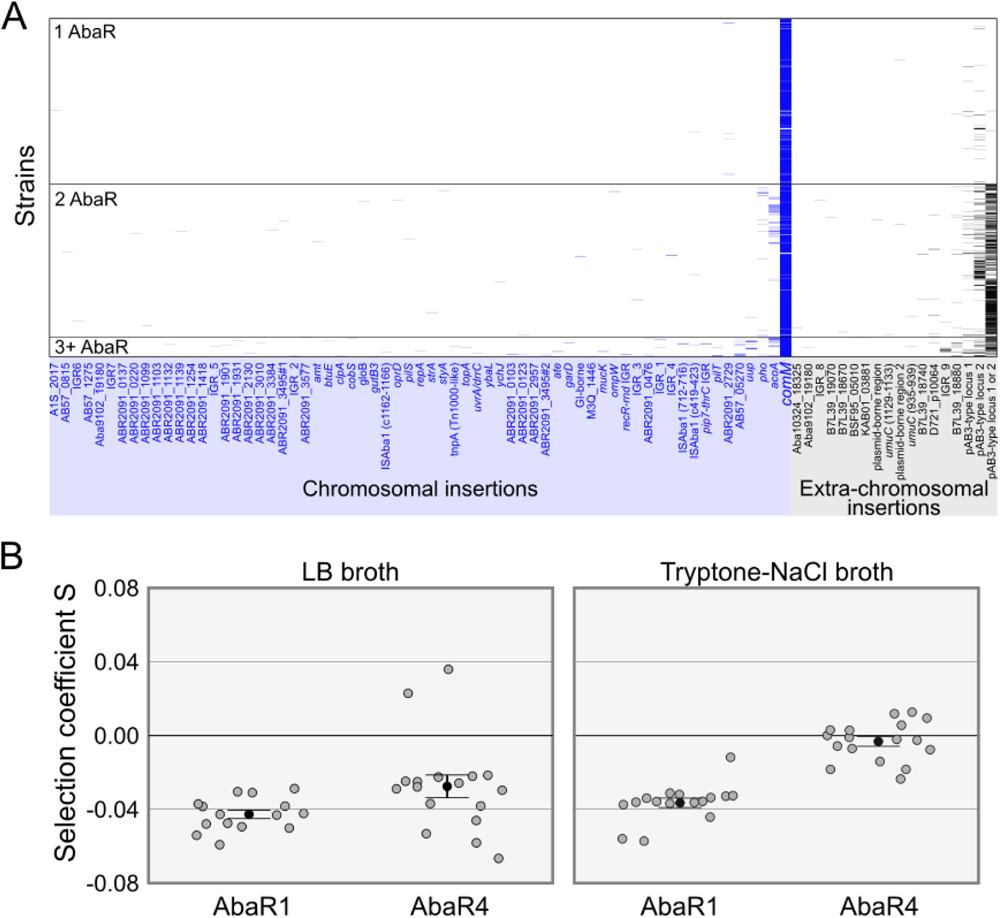
AbaR are preferentially inserted in the *comM* gene and impose a fitness cost. A. Genomic distribution of AbaR islands in *A. baumannii* genomes. Insertion sites are listed on the x axis. Strains are sorted on the y axis by the number of AbaR insertions. Data were retrieved from work by Bi et al., 2019. Names of some insertions site were abbreviated (see Material and methods). B. Experimental determination of the cost on fitness of AbaR4 and AbaR1. The effect on fitness of acquisition of AbaR4 and AbaR1 by the M2 strain were measured using bacterial competitions with their parental M2 strain in LB medium or in tryptone-NaCl medium. Selection coefficient (S) of the AbaR-carrying strains were determined relative to the parental M2 strain. Independent measures of selection coefficients are represented by the dotplot, black dots represent mean values and error bars represent the standard error.

### AbaR impose a fitness cost on their host

Acquisition of a single AbaR confers host with resistance to multiple antibiotics (47), providing a strong selective advantage under potentially diverse conditions of antibiotic stress. However, fixation of MGEs in bacterial genomes depends on their fitness effect on the bacterial host in absence of antibiotics (48). We therefore assessed the cost of AbaR acquisition by a naive host, *A. nosocomialis* strain M2, using an isogenic bacterial competition assay. We selected two distinct AbaRs, AbaR1 and AbaR4. AbaR4 is an emerging, 16-kbp long AbaR, that contains the Tn*2006*-*bla*_OXA-23_ transposon conferring resistance to carbapenems (49–52). In contrast, AbaR1 is an 86-Kb long AbaR including 25 putative resistance genes (53), conferring resistance to several old-generation antibiotics (aminoglycosides, cephalosporins, tetracycline, and sulfonamide) (47), and which is not found in modern isolates. AbaR1 and AbaR4 are found inserted in the *comM* gene in the chromosome of strains AYE and 40288, respectively. To disentangle the effect on fitness of the insertion site and of the island, we first quantified the fitness cost of the inactivation of *comM* by running competitions between M2 and M2 Δ*comM* strains, alternatively labeled by the introduction of the Hu-sfGFP marker (marker swap to deduce its cost on fitness, see Material and Methods). The selection coefficient of the M2 Δ*comM*, corrected from marker cost, appeared weak and negative (mean=-0.011, SE=0.005). Thus, inactivation of *comM* does not confer a fitness advantage that could have simply explained the high occurrence of AbaRs at this site. Next, to assess the effect on fitness of the selected AbaR, they were horizontally transferred to the M2 recipient to perform competition experiments between parental and AbaR-carrying derived strains. Competitions were conducted in rich and low-nutrient media (LB broth and tryptone-NaCl media, respectively). Acquisition of AbaR1 was deleterious to their host cells with selection coefficients of -0.043 and -0.037 in LB and tryptone-NaCL, respectively (Figure 1B). The acquisition of AbaR4 seems less deleterious than that of AbaR1, with a selection coefficient of -0.027 in rich LB medium. Similarly to AbaR1, this cost is lower in the tryptone-NaCl condition. In both media, AbaR1 showed a higher fitness cost than AbaR4. This could be due to its large size (86 kb), and is consistent with the higher prevalence of shorter AbaR in *A. baumannii* genomes (10-20 kb) (11). Overall, the data suggests that without selective pressure, AbaRs tends to incur a fitness cost. However, these fitness costs can be influenced by factors such as the content and size of the islands, as well as the prevailing growth conditions.

### Within-population natural transformation favors deletion of AbaR

Transfer of AbaR from AbaR-infested individuals to uninfested individuals spontaneously occurs in mixed populations (24). These within-population HGT are attributed to the natural release of DNA by some cells, which has the capacity to transform other cells within the population. Genomic analyses showed that within-population acquisition of AbaR results from the import of large, two-digits kb-long DNA fragments (24). Such fragments, naturally released by AbaR-free cells could also promote the deletion of the island in AbaR-infested cells.

We thus designed a genetic setup to determine the rate at which bacteria acquire or delete AbaR of variable length from 2 to 16 Kb, and so in realistic populations. To do so, we constructed a set of mutants, called M2 AbaR4-*Lj* and which carry an AbaR4-derived insert (*j*) of variable length (*L*) (Supplementary Figure S1A). Another set of mutants, called M2 AbaR4-Δ*Lj* was produced by deleting the insert while keeping the flanking regions (Supplementary Figure S1A). The pairwise combination of these two sets of mutants in a mixed population makes it possible to measure the rate at which the AbaR4-derived insert is acquired or deleted. Indeed, AbaR4-ᐃ*Lj* cells which acquired the insert using DNA released by AbaR-*Lj* could be detected by the gain of the kanamycin resistance conferred by the *aphA* gene (Supplementary Figure S2A). In contrast, AbaR-*Lj* cells which took up DNA released by AbaR4-ᐃ*Lj* cells will have the insert deleted, reconstituting the *aacC4* gene conferring resistance to apramycin (Supplementary Figure S2B). In each population consisting of a mix of M2 AbaR4-*Lj* and M2 AbaR4-Δ*Lj*, either strain could act as donor or recipient. To distinguish between these events, we alternatively deleted the *comEC* gene in one genotype to make it a donor-only (transformation is inactivated by loss of *comEC*) while introducing the selectable *rpoB*(rif^R^) allele conferring resistance to rifampicin in the recipient genotype.

We observed that the acquisition rate of the insert decreases with its length (Supplementary Figure S2C), confirming previous results that the full-length, 16.6 kb-long AbaR4 island is acquired 4-times less efficiently than the 4.8 kb-long Tn*2006* (24). Yet, while statistically significant (Pearson correlation test; n=101, t=-2.467, p-value=0.0153), the size effect remained limited with a slope of - 2.7×10^−2^ kb^-1^. In contrast, deletion rates do not vary significantly with the length of the insert (Pearson correlation test; n=69, t=1.865, p=0.066) indicating that small and large islands are deleted with similar efficiencies (Supplementary Figure S2D). Importantly, in the range of 10-16 kb, deletion is more efficient than acquisition (Figure 2A) (Student’s T test; df=84, t=-4.89, p=4.76 E-6), an asymmetry that is in line with previous results (54). This indicates that, in the absence of selective pressure to maintain AbaR, natural transformation would be effective at purging them from the population.

**Figure 2.**
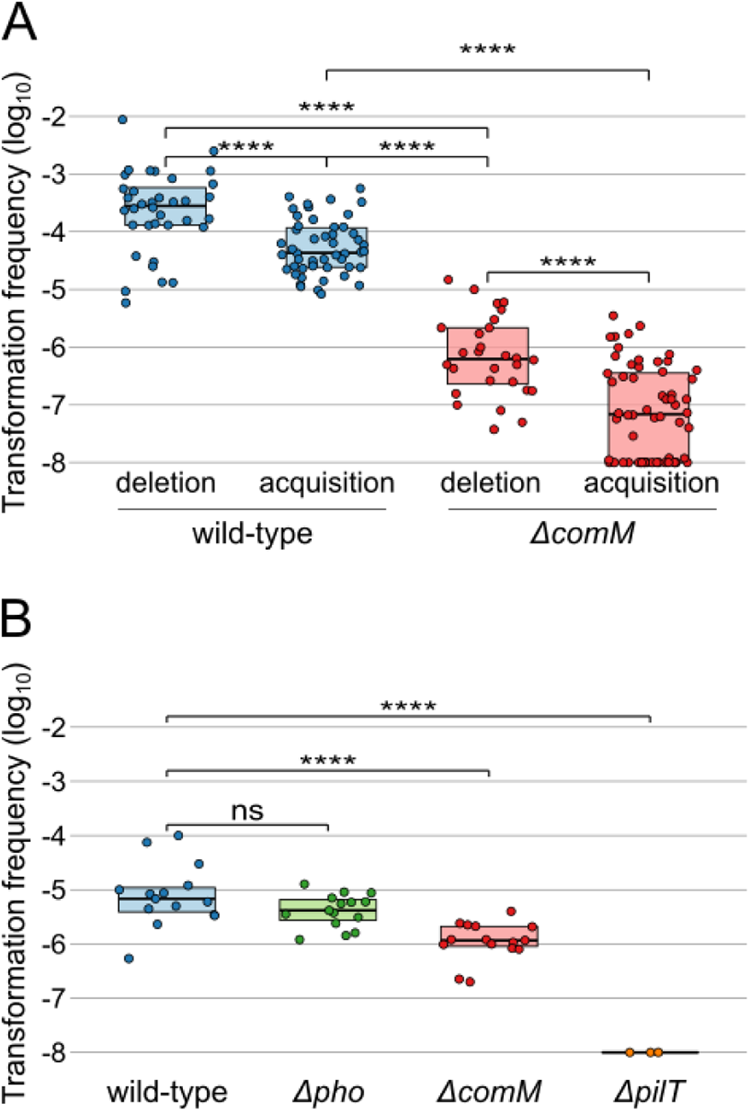
Frequency of natural transformation events spontaneously occurring within mixed populations. A. Frequency of transformation events resulting in the deletion or acquisition of inserts of length from 10 to 16 kb in a wild-type strain and Δ*comM* mutant. Equal parts of strains carrying or not the insert are mixed. Following incubation, the frequency of cells not previously carrying the insert and which acquired the insert is determined (acquisition). Similarly, the frequency of cells which have lost the insert (deletion) is determined (see Methods). B. Frequency of transformation events resulting in the acquisition of a SNP allele conferring resistance to rifampicin. As in A, equal parts of strains carrying or not the allele are mixed. The fraction of cells which acquired the SNP from the other cells is determined by plating (see Methods). (ns:p>0.05; *:p≤0.05; **:p≤0.01; ***:p≤0.001; ****:p≤0.0001). The detection limit of transformation events is 1×10-8. Boxplots represent the 25th percentile, median, 75th percentile.

### Within-population natural transformation events are differentially altered by *comM* inactivation

Depending on their insertion site, the acquisition of AbaR could alter the rate of subsequent within-population gene and allelic transfer events. Indeed, *comM*, encodes a DNA helicase implicated in homologous recombination of transforming DNA. When bacteria are artificially provided with exogenous DNA carrying a single nucleotide polymorphism (SNP), the loss of the ComM helicase reduces the transformation frequency by ~7-fold in *Acinetobacter baylyi* (46). Using exogenously added DNA, we confirmed this effect in *A. nosocomialis*, with the deletion of *comM* causing a ~6-fold reduction in the rate acquisition of the SNP of the *rpoB*(rif^R^) allele conferring resistance to rifampicin (Supplementary Figure S2E). In contrast, inactivation of *pho*, a gene of unknown function and in which AbaR is also found, has no effect on the rate of allelic transfer conferring rifampicin resistance. In contrast, inactivation of *pilT*, encoding the retraction ATPase of Type IV pilus required for transformation, completely abolished allelic transfer conferring rifampicin resistance (Supplementary Figure S2E).

We next tested the impact of *comM* inactivation on allelic transfer spontaneously occurring in the mixed-population setup, which here consisted in equal parts of the M2 cells carrying or not the *rpoB*(rif^R^) allele. To detect the acquisition by rifampicin-sensitive recipient M2 cells, we used strains carrying a kanamycin resistance marker at a neutral site (*att*Tn*7*) in order to distinguish them from *rpoB*(rif^R^) allele donor cells. Also, to avoid the confounding effect of the *rpoB*(rif^R^) allele donor cells acquiring the kanamycin resistance marker of recipients, the *comEC* gene of recipients was inactivated. We found that recombinants displaying both kanamycin and rifampicin resistance were produced at a frequency of 1.×10^−5^ (Figure 2B). Inactivation of *pho* had no effect. Confirming that HGT in these conditions are due to natural transformation, inactivation of *pilT* abrogated transfer of the *rpoB*(rif^R^) allele. Inactivation of *comM* resulted in a 8-fold reduction in the acquisition of the rifampicin resistance from neighboring cells (Figure 2B). Thus, the inter-individual transfer of SNP alleles is marginally affected by the inactivation of *comM*. Given the role of ComM in integration of heterologous DNA (46), we next tested the impact of *comM* inactivation on within-population acquisition and deletion of large heterologous DNA sequences. Bacteria lacking the *comM* gene displayed decreased acquisition rates of the heterologous insert *j* by around 2-logs (Figure 2A). Interestingly, the absence of functional ComM did not cause any additional defect in acquisition of inserts of increasing length with a slope of -6.9×10^−2^ kb^-1^, similar to that observed in wild-type cells (Supplementary Figure S1C). Deletion rates of inserts follow the same trend, being strongly reduced by the inactivation of *comM* with little dependence on the length of the deleted sequence (Supplementary Figure S1D). Like in the wild-type situation, deletion rates remain higher than acquisition rates. Importantly, deletion rates in *comM*-defective individuals are ~50 times lower than acquisition rates in wild-type cells (Student’s T test; df=76; t=14.382, p<2.2×10-16) (Figure 2A). This suggests that AbaR islands inactivating *comM* may not be purged by natural transformation within a population.

### Transformation-inhibiting AbaRs gain a competitive advantage in a fluctuating environment

The asymmetry of natural transformation supports a role in removing parasitic MGEs from the chromosome of bacteria (genome curing effect). It could be hypothezised that insertional inactivation of the *comM* gene would represent an evolutionary response of AbaRs. We found that AbaR insertion in *comM* globally inhibits the transformation rate of bacteria. However, the extent of this inhibition appears heterogeneous between DNA acquisition and DNA removal and between large DNA fragments (gene transfer events) and SNPs (allelic transfer). To disentangle the effect of *comM* inactivation, we developed a computational model to follow the dynamic of AbaRs within a simulated bacterial population (See Material and Methods and Supplementary Figure S3A). We considered three competing AbaRs with different insertion sites. AbaR inserting in *comM* (*comM*::AbaR) which partially inhibits transformation of their host, AbaR inserting in *pilT* (*pilT*::AbaR) representing AbaR which completely prevents transformation, and AbaR inserting in *pho* (*pho*::AbaR) considered as a generic neutral site with no incidence on natural transformation. Simulated bacterial cells are allowed to divide following a logistic growth model. Cells can thus naturally die at a basal rate, and at an elevated rate when encountering environmental stress. Consequently, their DNA is released, becoming available for other organisms to undergo natural transformation. The capacity for natural transformation of individuals is dictated by their genotype (wild-type, *pilT, pho, comM*) and calibrated from empirical data (Figure 1 and 2, summarized in Supplementary Table S1 and S2). AbaR could confer resistance to antibiotic stress but would reduce the growth rate of their host cell (fitness cost) according to experimental values (Figure 1B).

To examine the impact of transformation asymmetry in MGE removal, we initially observed the dynamics of AbaR under an hypothetical scenario where they incur no costs on fitness and do not provide any selective advantage to their bacterial host. We found that when bacteria carrying AbaR at the neutral site (*pho*::AbaR) are equally mixed with uninfested wild-type cells, AbaR are rapidly purged from the population (Supplementary Figure S3B). Thus, the transformation asymmetry in our system of within-population transformation is effective to cure a genomic island (here AbaR) from a population (Supplementary Figure S3B). In contrast, *comM and pilT-*inactivating AbaRs are not purged and even spread to infest the entire population, thereby confirming the genome-curing function of natural transformation (Supplementary Figure S3B). Furthermore, still without incurring costs or facing antibiotic-induced stress, when all four genotypes are mixed in the initial population, transformation-inhibiting AbaRs (*comM*::AbaR or *pilT*::AbaR) coexist quasi-neutrally and outcompete the AbaR that does not (*pho*::AbaR) (Figure 3A). We then considered the more realistic situation where AbaRs are costly and confer antibiotic resistance. In the absence of stress, wild-type cells expectedly gain a competitive advantage over any AbaR-carrying cells (Supplementary Figure S3B). Single occurrence of stress then strongly favors AbaR-carrying cells, which take over the population under constant or stochastic stress (Supplementary Figure S3B). In simulations initialized with only the wild-type genotype, while AbaRs are added to the extracellular compartment at a residual rate, short and rare exposures to antibiotic stress are sufficient to select and maintain AbaRs (Figure 3B and 3C). If stress is scarce (mean frequency f=1e-4) and short (mean duration=100), the wild-type genotype is favored (Figure 3D, with a low proportion of cells carrying AbaRs (Figure 3C). In this situation, all AbaR-carrying genotypes perform equally well. In contrast, under constant stress (Supplementary Figure S3A) or frequent and/or longer stress duration, which increase the proportion of AbaR-carrying cells (Figure 3C), *pho*::AbaR appear less competitive and can even become extinct, while the transformation-inhibiting AbaRs (*comM* or *pilT*::AbaR) coexist in a quasi-neutral way. This stems from the genome curing effect: the inevitable removal of AbaR from the transformation-proficient *pho*::AbaR cells, while transformation-inhibiting AbaR (*comM* and *pilT*) are themselves protected from this effect.

**Figure 3:**
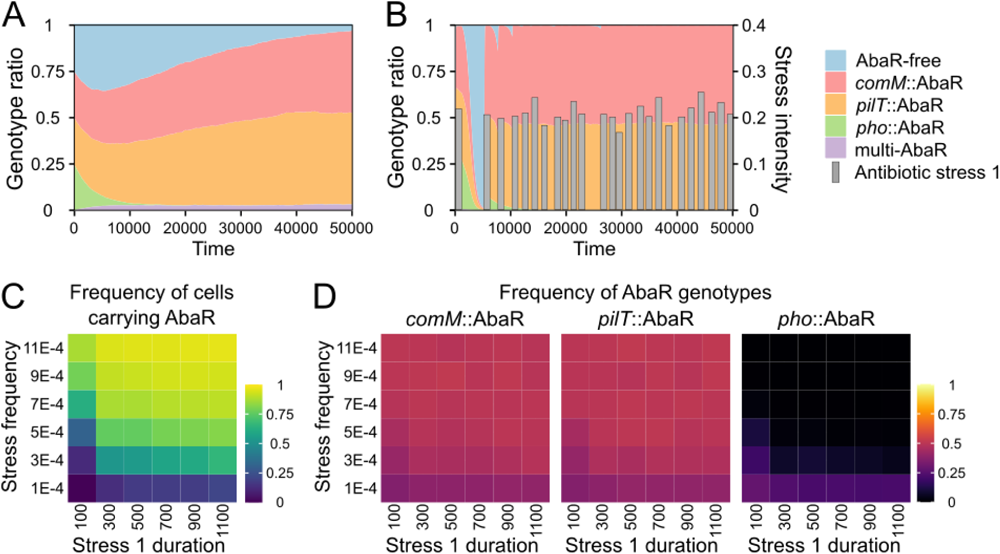
Transformation-inhibiting AbaRs gain a competitive advantage in a fluctuating environment. Temporal dynamics of bacterial genotypes carrying hypothetically costless AbaR in the absence of antibiotic stress (A) or in presence of a stochastic stress and AbaR imposing a fitness cost (0.01) (B) (mean peak duration=1.100, mean peak frequency=1.1E-3, mean peak intensity=0.2). C. Mean frequency of cells carrying AbaR between times 30.000 and 50.000 as a function of stress duration and stress frequency (averaged from 100 simulations each). D. Mean frequencies of each AbaR genotype between times 30.000 and 50.000 generations as a function of stress duration (*d*) and stress frequency (*f*) (averaged from 100 simulations each). Stress duration (*d*) is expressed in time steps, with 1 generation corresponding to about 3 time steps. Model parameters are listed in Supplementary Table S1 and S2.

Altogether, these findings demonstrate the potency of transformation asymmetry in purging the genome of AbaRs and other MGEs. When AbaR provides a selective advantage during stochastic exposure to antibiotic stress, the AbaR variants that hinder transformation gain a competitive advantage over those that do not. Interestingly, *comM*::AbaR and *pilT*::AbaR genotypes show similar dynamics, suggesting that a moderate inhibition of natural transformation is nearly as effective as a complete loss of natural transformability.

### Transformation-modulating AbaRs targeting *comM* maintain the adaptability of their hosts

A role of natural transformation in removing parasitic MGEs remains compatible with its involvement in adaptability through recombination of beneficial alleles (26). The high rates of recombination in *A. baumannii* indicate a major role of natural transformation in acquisition of beneficial alleles. This is exemplified by the acquisition of the highly selective SNPs in *gyrA* and *parC* genes conferring resistance to fluoroquinolones (18). We then implemented allelic variations in the computational model by introducing a SNP providing resistance to a specific stress 2. Stress 1 to which AbaR are conferring resistance and stress 2 occur stochastically and independently.

When stress 2 is rare and the cost of carrying the SNP is low (Figure 4A), as previously, *pho*::AbaR, which cannot oppose the genome curing effect, is eliminated in favor of AbaRs which inhibit transformation. However, here we detect a clear advantage of *comM*::AbaR over *pilT*::AbaR. *comM*::AbaR, which maintains a high level of transformation for short sequence changes, allows their host bacteria to acquire the inexpensive SNP that remain relatively abundant in the population (Supplementary Figure S4). In contrast, *pilT*::AbaR that completely inhibits transformation cannot respond as rapidly to changes in exposure to stress 2 and are at a disadvantage. It should be noted that *pilT*::AbaR are not completely counter-selected because wild-type cells (carrying or not the SNP), which grow during stress 1-free periods, can subsequently acquire *pilT*::AbaR by transformation, resulting in the persistence of *pilT*::AbaR in the population (Supplementary Figure S4). When Stress 2 is rare but the cost of carrying the SNP is very high (Figure 4B), the SNP becomes extremely rare in the population (present only at a residual frequency, Supplementary Figure S4) and transformation becomes ineffective in relation to the stress 2. The results of the simulations then mirror those without stress 2 (Figure 3D) in which *comM*::AbaR and *pilT*::AbaR coexist neutrally while *pho*::AbaR is cleaned up by transformation.

**Figure 4:**
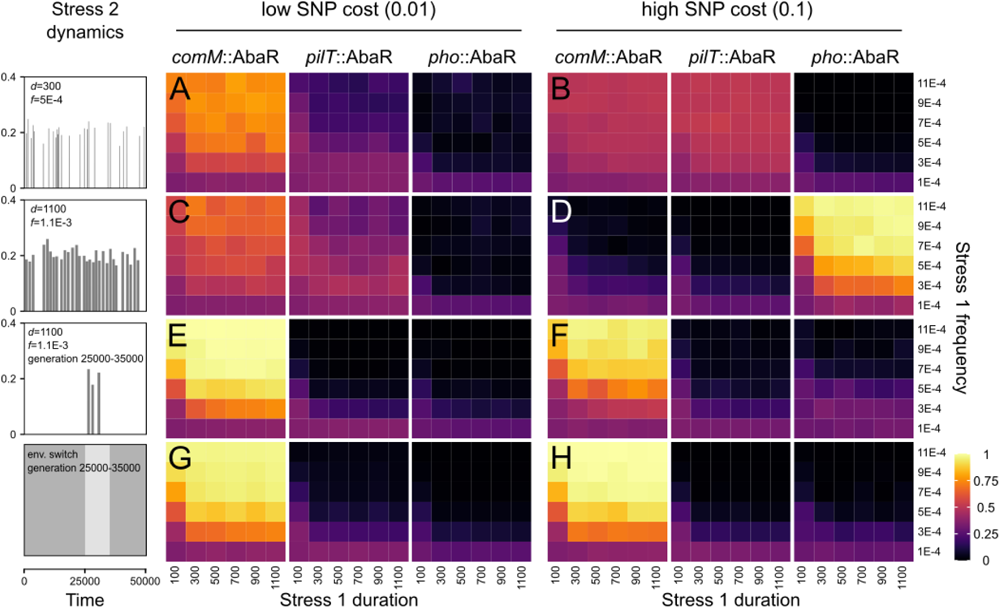
Frequency of AbaR islands in environments favoring adaptation by allelic transfer. Temporal dynamics of bacterial genotypes carrying AbaR in environments exposed to a stochastic stress 1 (to which resistance is conferred by AbaR) and another environmental change 2 to which advantage is conferred by a SNP acquired by allelic transfer. Grey boxes show the dynamics of stochastically occurring stress 2, which from top to bottom are rare and short (mean peak duration *d*=300, mean peak frequency *f*=5E-4), frequent and long (mean peak duration *d*=1100, mean peak frequency *f*=1.1E-3), limited to a short period (25000-35000 time units). Lastly, environmental change 2 is not a stress but a change in resource availabilty occurring between 25000 and 35000 time units. The ratio of bacterial genotypes carrying AbaR unders stress 2 environments were assessed as a function of stress 1 frequency and duration (panel A to H). The cost on fitness of the SNP conferring resistance to stress 2 was set as low (0.01, identical to the fitness cost of AbaR) (panel A, C, E, G) or high (0.1) (panel B, D, F, H). Mean frequencies of each AbaR genotype between times 30000 and 50000 as a function of stress duration and stress frequency (averaged from 100 simulations each). Model parameters are listed in Supplementary Table S1 and S2.

When stress 2 is frequent and the SNP is inexpensive (Figure 4C), the results and their explanations are similar to the situation when stress 2 is rare. On the other hand, when the SNP is very costly and exposure to stress 2 is frequent (Figure 4D), the SNP persists in the population and maximum transformation activity is selectively advantageous: this ensures great genomic plasticity, i.e. acquisition of the SNP when exposed to stress 2, but also its rapid elimination when it is no longer needed. Bacteria carrying *pho*::AbaR can acquire and eliminate SNPs 8-times faster than bacteria carrying *comM*::AbaR. As a result, *pho*::AbaR has a clear selective advantage and becomes fixed in many contexts of stress 1 exposure. Note that *pho*::AbaR does not escape genome curing but as it becomes the only AbaR capable of conferring stress 1 resistance, it is regularly recruited/reintegrated by transformation. These results could be interpreted as a form of cooperation (mutualism) between *pho*::AbaR and their bacterial host, but this does not seem robust from an evolutionary point of view (see below).

We then tested situations in which the environmental change occurs over a relatively short period during with the modeled trajectories (Figure 4E to 4H). This change is characterized either by exposure to a stress 2 to which the SNP confers resistance (Figure 4E and 4F), or by a change in the continuously available resource, which will be better exploited by the bacteria carrying the SNP (Figure 4G and 4H). In all these situations, the insertion in *comM* clearly confers a strong selective advantage. During long periods without stress 2, *pho*::AbaR is cleaned out in favor of *comM*::AbaR and *pilT*::AbaR (situation similar to Figure 2). As soon as environmental change occurs, *pilT*::AbaR is brutally counter-selected because, unlike *comM*::AbaR, it cannot acquire the SNP by transformation. The success of *comM*::AbaR is therefore explained by the fact that it maximizes its probability of persistence (reducing the probability of being eliminated by transformation to almost 0) while maintaining a non-negligible level of transformation activity for allelic changes allowing the host bacterium to respond quickly to the change in its environment. It should be noted that *pilT*::AbaR could persis if wild-type cells which managed to acquire the SNP subsequently acquire *pilT*::AbaR during the short environmental change. However, the newly formed *pilT*::AbaR bacteria would also be rapidly counter-selected once the initial environment was restored, in favor of *comM*::AbaR bacteria that were able to eliminate the SNP by transformation.

In conclusion, we found that if any allelic change is necessary for survival or adaptation to a reversible environmental shift, then even a single occurrence of such is sufficient to fix *comM*::AbaR in the population.

## Discussion

We investigated the insertion sites of AbaRs in *A. baumannii* genomes, quantified transformation rates in experimental populations, and performed modeling work to understand the insertion strategy of AbaR islands into the chromosome of bacteria. We found that insertion into the *comM* gene confers a selective advantage in many contexts, surpassing other insertion strategies that either do not inhibit (such as insertion into the *pho* gene) or completely inhibit transformation (such as insertion into the *pilT* gene). This *comM* insertion strategy should be selected due to two combined effects on transformation. First, inserting in *comM* allows AbaR to circumvent the genome curing effect of transformation. The genome curing effect was proposed on the basis of the transformation properties of *Streptococcus pneumoniae*, and requires that deletion rates be greater than acquisition rate (referred to asymmetry of transformation) (27). Such strong asymmetry was verified in *S. pneumoniae*, with experimental models involving the addition of exogenous DNA (54). We here confirm this phenomenon in *A. baumannii*, yet in a unique and more realistic model in which transformation occurs spontaneously within a population, not through the addition of exogenous DNA. In this system, the degree of transformation asymmetry is less pronounced than in the *S. pneumoniae* model. This is caused by the rates of acquiring naturally-released DNA which are far less affected by the size of the acquired DNA in our model (2.7×10^−2^ kb^-1^) than in the *S. pneumoniae* model (1.08×10^−6^ kb^-1^). This is possibly due to the fact that naturally released DNA present fewer transformation-terminating nicks than purified DNA. Simulations indicate that even a limited asymmetry is effective to cure AbaR - or any other chromosome-integrated MGE - from a population (Supplementary Figure S3). Through the inactivation of *comM*, AbaR significantly diminish the deletion rate of their host, leading to a substantial 50-fold asymmetry between the rate at which wild-type cells can acquire AbaR and the rate at which they can subsequently remove it. By counteracting the curing effect of natural transformation, AbaR inserting in the *comM* gene can invade a population (Supplementary Figure S3).

Second, the success of AbaR is explained by the fact that inserting in *comM* differentially inhibit allelic and gene transfer events. Inactivating *comM* only marginally affects the transfer of SNP by transformation. While AbaRs inserting in *comM* can oppose the genome curing effect of transformation, their host cell would still allow the transformation-mediated recombination events which can shuffle mutations, break linkage desequilibrium, acquire beneficial mutations or purge deleterious ones. We observed that AbaR inserting in *comM* outcompete AbaR inserting in any other site, whether allelic transfer confers a strong selective advantage (resistance to antibiotic) or a milder adaptive advantage linked to the exploitation of a new resource in the environment (Figure 4). This differential inhibition of allelic and gene transfer events is linked to the role of ComM as the helicase dedicated to natural transformation. Inactivation of any other genes of the transformation pathway, and involved in initial binding (type IV pilus genes, *pilS, pilT, pilA*, etc), import (*comEA, comEC, comFC*), protection (*dprA*) or recombination (*recA*) of DNA would not result in such differential inhibition (23). Thus, *comM* represents a unique target to block genome curing while still allowing evolution by acquisition and recombination of new alleles.

Our simulations demonstrate that the *comM* insertion strategy is advantageous when the model incorporates some degree of environmental complexity, such as different types of stress, or stress combined with changes in environmental resources. It would thus seem essential to consider the complexity of the environment to understand the prevalence of AbaR insertion into the *comM* gene of *A. baumannii*. Distinct MGEs have been found to disrupt the *comM* gene in *Mannheimia succiniciproducens* (55), *Mannheimia haemolytica (55), Francisella philomiragia, Pseudomonas syringae* (27) and *Aggregatibacter actinomycetemcomitans* (56). The latter two species are pathogens known to be naturally transformable. Hence inserting in *comM* may be a successful strategy for many other MGEs of populations/species evolving in contrasting environments. Generally, it would be necessary to rigorously describe MGE insertion strategies, characterize their consequences on natural transformation, and identify ecological contexts (e.g., hospital and non-hospital environments) exerting selective pressure that favor certain insertions over others.

The *comM* insertion strategy may benefit only certain types of MGEs. Total inhibition of transformation, which is not a successful strategy in our model, has been reported for integrative conjugative elements (ICEs) (39–41, 43) and expected for prophages inserting into genes essential for transformation (27). In contrast to genomic islands such as AbaRs, these types of MGEs are autonomous in their horizontal transfer and do not depend on natural transformation to access a new genetic context. Therefore, they can completely inhibit transformation while escaping extinction in the event of environmental change by controlling their own horizontal transfer. On the contrary, non-autonomous MGEs (such as AbaRs) depend on natural transformation to change their genetic context and may have an interest in maintaining basal transformation activity. Interactions between MGEs, should also be considered to understand the strategies of MGEs in inhibiting transformation. For example, autonomous MGEs could still benefit from the inhibition of transformation conferred by AbaRs inserted into the *comM* gene. These MGEs could then dispense with carrying transformation inhibition mechanisms.

The insertion strategy into the *comM* gene should be a key element in the dynamics of antibiotic resistance genes, at least in *A. baumannii*. This may occur in two distinct ways: first, AbaRs inserted in *comM* become the playground for genetic innovation, as they persist for a long time in bacteria genomes and can accumulate transposons or undergo rearrangements, easily leading to the evolution of accessory functions such as antibiotic resistance. Second, AbaRs can have their own intracellular dynamics and change vehicles (genetic support). AbaRs, which may have conserved a transposition activity, can jump to plasmids, thus facilitating new pathways for the spread of antibiotic resistance and new evolutionary trajectories of AbaRs on these new genetic vehicules. In this sense, our genomic analysis reveals that when an AbaR is present on a plasmid, it is also almost always present in *comM* - although the reverse is not true -, suggesting intracellular movements (Figure 1A).

In conclusion, studying the strategies of MGEs and their interactions with the natural transformation mechanism of bacteria is crucial for understanding the persistence and evolution of antibiotic resistance. Our findings highlight the importance of considering environmental complexity when investigating the prevalence of specific transformation inhibition strategies, such as the insertion into the *comM* gene. By uncovering the mechanisms explaining the successful establishment and long-term persistence of MGEs, we could better develop strategies to combat the spread of antibiotic resistance and preserve the effectiveness of antimicrobial interventions.

## Materials and methods

### AbaRs insertion sites in bacterial genomes

AbaR-type insertion sites identified from 3,148 genomes were retrieved from Bi et al (11). Insertion sites belonging to the same gene (or intergenic region, IGR) were combined. Each insertion site is attributed to a chromosomal or extra-chromosomal category. AbaRs inserted into tetA sites, which correspond to an AbaR inserted within AbaR were removed from the data. AbaR-type insertions were detected by the presence of either the conserved sequence left or right (CSL, CSR) of AbaR-type elements (11). For the AbaRs inserted in the pAB3-type locus 1 and pAB3-type locus 2 sites, we checked that both boundaries were present. Finally, we represent the insertion sites of the different AbaRs for each bacterial strain, by grouping the strains according to the number of AbaRs they carried (analysis and graphical representation: R4.3.1 and ggplot2). The name of some insertion sites were abbreviated as follows: IGR_1, ABR2091_3231-tRNAArg; IGR_2, ABR2091_3550-ABR2091_3551; IGR_3, ABR2091_0006-ABR2091_0007; IGR_4, CBI29_04511-CBI29_04512; IGR_5, ABR2091_1794-ABR2091_1795; IGR_6, AB57_0986-AB57_0987; IGR_7, ABR2091_0111-ABR2091_0112; IGR_8, ACX61_19590-ACX61_19595; IGR_9, CBI29_04511-CBI29_04512

### Bacterial strains, plasmids, and genetic constructs

Bacterial strains with genotypes and plasmids are detailed in Supplementary Table S3. All the oligonucleotides used in this study for genetic modification are listed in Supplementary Table S4.

### Chromosomal modifications in *Acinetobacter*

Genetic constructions were performed using overlap extension PCR to create assembly PCR fragments carrying the genetic construction with a selection marker flanked by 2 kbp-long fragments that are homologous to the insertion site. A scarless genetic strategy was used to create the mutations without antibiotic resistance markers (47). All PCRs for genetic modifications were performed with a high-fidelity DNA polymerase (PrimeStarMax, Takara) and chromosomal modifications were confirmed by colony PCR (Dream Taq, Thermo Fisher).

### Bacterial strains for fitness cost determination

The M2 *hu*-*sfgfp aacC4* (Hu-sfGFP) strain was obtained from a previous work (21). The M2 *comM*::AbaR4 and M2 *comM*::AbaR1 strains were generated by naturally transforming the M2 *comM*::*sacB aacC4* with genomic DNA from *A. baumannii* strains 40288 (AbaR4) (24) AYE (AbaR1) (53). Fluorescent variants were generated by naturally transforming bacteria with a PCR product of the *hu*-*sfgfp aacC4* construct.

### Bacterial strains for SNP acquisition

The M2 *rpoB*(Rif^R^) rifampicin resistant variant of the *A. nosocomialis* M2 wild-type (WT) strain was obtained by plating an overnight bacterial culture on a LB agar plate supplemented with rifampicin. Spontaneous resistant mutants were then purified and the *rpoB* gene from each one was further sequenced. The selected isolate displayed normal growth on LB plates supplemented with rifampicin and the non-synonymous mutation A1565T (G522L) in the *rpoB* gene. The non-transformable derivative M2 *rpoB*(Rif^R^) *comEC*::*aacC4* strain was obtained by naturally transforming the M2 *rpoB*(Rif^R^) strain with a PCR product of the *comEC*::*aacC4* construct. The M2 Δ*comM* strain has a deletion of 231 bp in the coding sequence of the *comM* gene and the M2 Δ*pho* strain has a deletion of 825 bp in the coding sequence of the *pho* gene. Both mutants were generated using the scarless genetic strategy. The M2 *pilT*::aphA was obtained from Gottfried Wilharm (20). To test acquisition of the SNP in mixed culture with the M2 *rpoB*(Rif^R^) *comEC*::*aacC4* strain, recipient strains were made kanamycin resistant by inserting the marker at the attn7 site.

### Bacterial strains for acquisition and deletion of inserts in mixed culture

*A. nosocomialis* M2 mutants carrying heterologous DNA chromosomal inserts with variable length were generated following the steps described in (Supplementary Figure S1B). Rifampicin resistant derivatives for selection of recipient strains were obtained by transforming bacteria with a PCR product of the *rpoB* gene amplified from genomic of the rifampicin resistant M2 *rpoB*(Rif^R^) strain. Non-transformable derivatives for donor strains were obtained by transforming bacteria with a PCR product of the *comEC*::*tetA* chromosomal modification.

### Fitness cost of AbaR

Bacterial competitions were performed by competing an *A. nosocomialis* M2 strain carrying an AbaR island inserted in the *comM* gene (M2 *comM*::AbaR4 or M2 *comM*::AbaR1 against the isogenic M2 with no AbaR. A single chromosomal fluorescent Hu-sfGFP marker was alternately expressed in each genetic background, as this marker discriminate fluorescent cells from the non-fluorescent cells using flow cytometry. The fitness cost associated with the expression of the Hu-sfGFP marker was determined by conducting marker swap experiments and subtracted from the fitness data points. Bacterial competitions were performed as previously described (57). Briefly, bacteria from the competing strains were grown in the competition medium (LB broth or Tryptone-NaCl medium) for one night. The next day, overnight cultures were mixed at equal ratio in a 24-wells plates and were pre-incubated at 37°C for 1 hour. Competitions were then performed in 12-wells plates by diluting 1000-fold each day the competing culture in a fresh liquid medium and incubated for 24 hours (10 generations per day). The ratio of AbaR carrying cells to wild-type cells was measured by counting 105 total cells using flow cytometry (Attune acoustic flow cytometer) at 0 and 10 generations. Selection coefficients were determined using the model s = [ln R(t)/R(0)]/t where R is the ratio of mutant to wild-type cells (58).

### Within-population transformation in mixed cultures

Mixed cultures were setup as previously described (24). Strains were grown overnight on LB agar plates, then inoculated and grown in 2 mL of LB until the cultures reach an optical density (OD, 600nm) of 1. The bacterial broth were then diluted to an OD of 0.01 in PBS. Then equal volumens of bacterial suspensions were mixed by pairs. The mixture (2.5 µL) was deposited on the surface of 1 mL of tryptone-NaCl medium solidified with 2% agarose D3 (Euromedex) poured in 2 mL microtubes or in wells of 24-well plates and incubated overnight at 37°C. Bacteria resuspended in PBS were plated on LB agar plates without antibiotics and LB agar plates containing with appropriate antibiotics. Recombinants frequencies were determined through calculation of the ratio of the number of CFUs on antibiotic selection to the total number of CFUs on plates without antibiotics.

### Transformation assays with purified DNA

Bacteria were grown approximately 5-6 hours in LB liquid medium at 37°C and were then diluted in PBS to obtain bacterial suspensions of 10^7^ cells/mL. Then an equal volume of the bacterial suspension and the DNA substrate solution (at 100 ng/µL) were mixed together and 2.5 µL of this bacteria/DNA mix were spotted on the surface of a 1 mL transformation medium (tryptone 5 g/L, NaCl 2.5 g/L, agarose type D3 2%) freshly prepared in a 2-mL Eppendorf tube. After 20h at 37°C, bacteria were harvested by adding 200 µL of PBS in the tube and vortexing. Transformants were selected after plating the bacteria in selective agar media containing various antibiotics (apramycin 30 µg/µL, kanamycin 15 µg/µL, rifampicin 100 µg/mL, tetracycline 15 µg/mL, nalidixic acid 60 µg/mL). Transformation frequencies were determined by calculating the ratio of the number of transformant CFUs (counted in selective media) to the total number of CFU (counted in non-selective media).

### Modelling the dynamics of AbaRs

We used a stochastic numeric model to generate the dynamics of AbaRs, which have different insertion strategies and compete in the exploitation of a bacterial population. The model comprises two compartments, one composed of bacterial cells, the other of extracellular DNA containing wild-type alleles and AbaRs carrying resistance. The bacterial cells have three AbaR insertion site in their chromosome (*comM, pilT, pho*) which can be occupied by wild-type allele (*comM*::WT, *pilT*::WT, *pho*::WT) or AbaR (*comM*::AbaR, *pilT*::AbaR, *pho*::AbaR). Considering all combinations of DNA type, there is a total of 2^3^=8 possible genotypes *i*.

Bacterial population growth follows a logistic model and the number of replicating cells per genotype *I* and per time step *dt*, is determined using a binomial distribution:

*G*_*i,t*+*dt*_ ∼*Bin* (*μ*_*i,t*_×*dt, N* _*i,t*_) where *μ*_*i, t*_ is the replication rate (see supplementary materials for details of its calculation) and *N*_*i,t*_ is the total number of cells with genotype *i* at time *t*. Cells carrying an AbaR have a replication rate lower that that of WT bacteria.

Bacteria release DNA upon cell lysis, the number of lysed cells per genotype *i* and time step *dt* also follows a binomial distribution:

*L*_*i,t*+*dt*_ ∼*Bin* (*k*_*i, t*_×*dt, N* _*i,t*_) where *k*_*i, t*_ is the lysis rate per time step at time *t*. Bacterial populations are faced with stochastic stresses of random duration, frequency and intensity. In the absence of stress, cells are lysed at a basal rate and under stress exposure, the lysis rate of WT cells increases but remains unchanged for cells with an AbaR carrying resistance.

Bacteria can uptake eDNA from the extracellular compartment and replace their own using transformation, the number of competent cells undergoing a transformation event during a time step *dt* is determined using a binomial distribution:

*C*_*i, t*+*dt*_∼*Bin*(*T*_*i,t*_×*dt, N* _*i,t*_) where *T*_*i, t*_ is the transformation rate of genotype *i* at time *t* (see supplementary materials for details of its calculation).

AbaRs inserted into *comM, pilT* and *pho* inhibit transformation partially, totally or not at all, respectively (Figure 2). For each run, the transformation probability values were randomly drawn from experimental data (see Supplementary materials).

In the extracellular compartment, eDNA is degraded at a constant rate *R*_*j*_, the number of degraded eDNA molecules *j* per time step is determined using a binomial distribution:

*D*_*j, t*+*dt*_∼*Bin*(*R* _*j*_× *dt, A* _*j, t*_) where *A* _*j, t*_ corresponds to the number of eDNA of type *j*.

The extracellular compartment is supplied by eDNA from lysed cells, each lysed cell releasing DNA molecules corresponding to their DNA composition. In addition, eDNA is added at a marginal rate simulating residual arrival from neighboring populations (open system).

Environmental fluctuations only concern exposure to stress 1, for which resistance is conferred by AbaR (Figure 3). More complex environmental fluctuations were simulated by either introducing a new stress or by modelling a change in resource (Figure 4). In the latter two cases, adaptation to the new environment is conferred by a new SNP that can be acquired by transformation. Because of the new SNP insertion site, the number of genotypes is increased, and becomes 2^4^=16. Bacteria can then acquire or remove this SNP by natural transformation, and these DNA molecules are also present in the extracellular compartment. The previous equations remain true. (see supplementary materials for more details about stress modelling).

The model was implemented in C++ (gcc version 9.4.0), using Code::Blocks IDE (20.03).

### Statistical analysis and figures

All the statistical analyses and figures were produced on R4.3.1 (additional packages: ggplot2, forcats, ggpubr, scales).

## Acknowledgments

This work was supported by the LABEX ECOFECT (ANR-11-LABX-0048) of Université de Lyon, within the program “Investissements d’Avenir” (ANR-11-IDEX-0007) operated by the French National Research Agency (ANR) and by the RESPOND program of the Université de Lyon (UDL). ASG was also supported by VetAgro Sup (PhD funding). This project has received financial support from the CNRS through the 80|Prime program. This work was performed using the computing facilities of the CC LBBE/PRABI.

