## Supplementary Materials for "Manipulation of natural transformation by AbaR-type islands promotes fixation of antibiotic resistance in *Acinetobacter baumannii* populations"

Computation model
Supplementary Table S1: Parameters used in the model.
Supplementary Table S2: Transformation event probabilities used in the model.
Supplementary Table S3: Bacterial strains and plasmids used in this study.
Supplementary Table S4: Primers used in this study.
Supplementary Figure S1. Strategy and mutant construction to quantify events of acquisition or deletion of a heterologous DNA fragment occuring within mixed populations.
Supplementary Figure S2. Quantification of rates gene transfer (acquisition and deletion) and rates allelic transfer (SNP) occuring within populations.
Supplementary Figure S3. A mathematical model of A. baumannii population involving HGT by natural transformation.
Supplementary Figure S4. Temporal dynamics and frequencies of bacterial genotypes carrying AbaR.

**Computational model**

We used a previous stochastic computational model (38), complemented with additional genetic elements and insertion sites in bacterial chromosomes. The model is composed of two compartments: bacterial cells (intracellular compartment) and extracellular DNA (extracellular compartment). Bacteria possess several insertion sites on their chromosome which can be occupied by two types of DNA each.

In the first stage, the environmental fluctuations only concern exposure to stress 1 (which occurs randomly), for which resistance is conferred by AbaR (fig. 3). Insertion sites of AbaRs include three genes, *comM*, *pilT* and *pho*, and can be occupied by their respective wild-type allele (*comM::*WT, *pilT*::WT or *pho*::WT) or by their respective AbaR genomic island (*comM::*AbaR, *pilT*::AbaR or *pho*::AbaR). In accordance with experimental results, the frequency of natural transformation can be affected by the inactivation of these genes, i.e. the insertion of an AbaR (see Supplementary Table S1 and S2). This frequency is unchanged when the *pho* gene is inactivated, is reduced when the *comM* gene is inactivated and is dropped to 0 when the *pilT* gene is inactivated.

In the second stage, we model more complex environmental fluctuations. In addition to stress 1 (for which resistance is conferred by AbaR), we are introducing either a new stress that reduces the survival of non-resistant bacteria, or a change in resource for bacterial growth (bacteria that do not have the allele adapted to this new resource have a lower growth capacity). In the latter two cases, adaptation to the new environment is conferred by a new single nucleotide polymorphism (SNP) that can be acquired by transformation and that integrates into a new site. The additional site can be occupied by a wild type allele or by a new SNP. The inactivation of the *comM* gene only slightly inhibits the acquisition of SNPs by transformation, whereas inactivation of the *pilT* gene totally inhibits it.

Considering all combinations of DNA type and scenarios, there is a total of $2^3=8$ (stage 1) or $2^4=16$ possible genotypes $i$ (stage 2). Bacteria can uptake eDNA from the extracellular compartment and replace their own using transformation. Bacteria release DNA upon cell lysis which can be enhanced by a stochastic bactericidal stress.

<u>Model processes</u>

Bacterial population growth follows a logistic model and the number of replicating cells per genotype $i$ and per time step $dt$, $G_{i,t+dt}$ is determined using a binomial distribution:

$$G_{i,t+dt} \sim Bin\left(\mu_{i,t}.dt, N_{i,t}\right) \quad (1)$$

where $\mu_{i,t} = \left(\mu_{max} - \frac{\mu_{max} - k_b}{K} N_{tot,t}\right) * \left(1 - c_i\right)$. (2)

$N_{i,t}$ is the number of cells with the genotype $i$ at time $t$. $\mu_{i,t}$ is the replication rate of genotype $i$ at time $t$. $\mu_{max}$ is the maximal growth rate. $k_b$ is the constant basal lysis rate, independent of the presence of stress. $N_{tot,t}$ is the total number of cells in the population (considering all genotypes) at time $t$. $c_i$ is the cost, in terms of cell replication, induced by all DNA types possessed by the genotype $i$. $K$ is the carrying capacity. The number of lysed cells per time step $L_{i,t+dt}$ is calculated using a binomial distribution:

$$L_{i,t+dt} \sim Bin\left(k_{i,t}.dt, N_{i,t}\right) \quad (3)$$

where $k_{i,t}$ is the lysis rate of genotype $i$ at time $t$ (see *Stress modelisation*). The natural transformation mechanism is modeled as 2 successive processes. First the number of competent cells engaging in a transformation event during a time step $C_{i,t+dt}$ is determined using the following binomial distribution:

$$C_{i,t+dt} \sim Bin\left(T_{i,t}.dt, N_{i,t}\right) \quad (4)$$

$$T_{i,t}=T_{max,i} * \left(\frac{\alpha A_{tot,t}}{1+\alpha A_{tot,t}}\right) \quad (5)$$

where $T_{i,t}$ is the transformation rate at time $t$, $T_{max,i}$ is the maximal transformation rate (i.e. the expected rate of transformation in the absence of any limiting constraint: (i) when the eDNA is not limiting, (ii) transformation is not inhibited and (iii) the cell acquires a short sequence - either a WT allele or a SNP-) of the genotype $i$, $\alpha$ is the binding rate between cells and eDNA, $A_{tot,t}$ is the total number of eDNA at time $t$. Genotypes with *pilT::AbaR* cannot uptake eDNA, $T_{max,i}$ is therefore equal to zero. For all others genotypes, $T_{max,i}=1\text{E-}3$ . Second, for cells having initiated a transformation event, a random eDNA molecule is selected in the extracellular compartment, considering the proportion of the different eDNA molecules. Depending on the eDNA molecule selected (large AbaR or short sequence such as WT allele or SNP) and on the integrity of the *comM* gene , the probability (S) of the successful integration of the focal molecule varies (see Supplementary Table S2). *S* probabilities are calculated from the natural transformation frequencies obtained in  experiments (results shown in Figure 2) and take into account the effects of transformation inhibition (following integration of AbaRs into *comM* or *pilT*) and the effect of sequence size (long AbaR sequence or short WT sequences or SNPs) originating from the extracellular compartment. To take account of errors in estimating transformation probabilities from experimental designs, for each run of the model, S were calculated from transformation probabilities drawn at random from the theoretical distribution of mean probability (distribution reported in Supplementary Table S2).

Cells undergoing a transformation event change their genotype according to the DNA type integrated. If the insertion fails, the eDNA is deleted and the genome of the cell remains intact. The overall variation of a genotype $i$ during a time step is summarized by:

$$N_{i,t+dt}=N_{i,t}+G_{i,t+dt}-L_{i,t+dt}-C_{i,t+dt}^{i\rightarrow !i}+C_{!i,t+dt}^{!i\rightarrow i} \quad (6)$$

where $C_{i,t+dt}^{i\rightarrow !i}$ is the number of competent cells going from the genotype $i$ to a different genotype and $C_{!i,t+dt}^{!i\rightarrow i}$ is the number of cells becoming of genotype $i$ from a different genotype.

In the extracellular compartment, eDNA is degraded at a constant rate $R_j$. The number of degraded eDNA molecules $j$ per time step, $D_{j,t+dt}$ is determined using a binomial distribution

$$D_{j,t+dt}\sim Bin\left(R_j . dt, A_{j,t}\right) \quad (7)$$

where $A_{j,t}$ corresponds to the number of eDNA of type $j$. The extracellular compartment is supplied by eDNA from lysed cells, each lysed cell releasing DNA molecules corresponding to their DNA composition. In addition, eDNA is added at a marginal rate simulating residual arrival from neighboring populations (open system). The number of eDNA molecules $j$ added per time step $M_j$ is defined by:

$$M_j=M_{input,j} * dt \quad (8)$$

where $M_{input,j}$ is the number of molecules of eDNA of type $j$ and $M_{input,j}$ is residual and set to be orders of magnitude lower than the DNA released by cell lysis. The overall variation of eDNA of type $j$ in the extracellular compartment during a time step is determined as follows by:

$$A_{j,t+dt}=A_{j,t}-D_{j,t+dt}+M_j+\sum_{i=1}^{n}\left[L_{i,t+dt}^{j}-C_{i,t+dt}^{j}\right] \quad (9)$$

where $L_{i,t+dt}^{j}$ is the number of cells of genotype $i$ (containing DNA of type $j$) which are lysed and $C_{i,t+dt}^{j}$ is the number of DNA molecules of type $j$ which are acquired by competent cells.

Stress modeling

AbaRs always confer resistance to stress 1, whereas the SNP can either confer resistance to stress 2 (Figure 4, lines 1-3) or confer a selective advantage when the bacterial population has access to a new resource over a given period (Figure 4, line 4).

If the AbaR and SNP confer antibiotic resistance, the lysis rate for genotype $i$ at time $t$ is calculated as follows:

$$k_{i,t}=k_b+I_{1,t} * \left(1-r_{1,i}\right)+I_{2,t}*\left(1-r_{2,i}\right) \qquad (10)$$

$I_{1,t}, I_{2,t}$ are the intensities of stresses 1 and 2 at time $t$, $r_{1,i}, r_{2,i}$ are the stress resistances provided by the DNA types possessed by the genotype $i$. Stress 1 increases the lysis rate of cells not carrying AbaRs whereas the lysis rate of cells carrying any AbaR remains at the basal rate $k_b$, the lysis rate of cells carrying the SNP remains at the basal rate in presence of stress 2 and the lysis rate of cells carrying both AbaR and SNP always remains at the basal rate.

Environmental fluctuations are modeled by two stochastic stresses whose durations and frequencies are variable. The probabilities that stress 1 or stress 2 start at each time are given by $N\left(f_1 \times dt, \sqrt{f_1 \times dt}\right)$ and $N\left(f_2 \times dt, \sqrt{f_2 \times dt}\right)$ where $f$ are mean stress frequencies. When a stress occurs, its duration is given by $N\left(d_1, \sqrt{d_1}\right)$ or $N\left(d_2, \sqrt{d_2}\right)$ where $d$ are mean stress durations, and the stress intensities $I_{1,t}$ and $I_{2,t}$ are given by $N\left(I_1, \sqrt{I_1}\right)$ and $N\left(I_2, \sqrt{I_2}\right)$ respectively where $I$ are mean stress intensities.

Under a scenario with a change of resource, we consider two SNP: $SNP_0$ and $SNP_1$. $SNP_0$ is advantageous in environment 0 and deleterious in environment 1 and inversely for $SNP_1$. The lysis rate for genotype $i$ at time $t$ is therefore calculated as follows:

$$k_{i,t}=k_b+I_{1,t} * \left(1-r_{1,i}\right) \qquad (11)$$

where $I_{1,t}$ is the intensity of stress 1 at time $t$, $r_{1,i}$ is the stress resistance provided by the genotype $i$, while the SNPs only influence the replication parameter (see equation (1)). The environmental switches are modeled as a sequence of environment 0 and 1.

#### Supplementary Table S1: Parameters used in the model.

| Parameter | Description | Values |
|---|---|---|
| $dt$ | Time step | 0.01 |
| $\mu_{max}$ | Maximal growth rate | 0.3 |
| $k_b$ | Constant basal lysis rate | 0.2 |
| $K$ | Carrying capacity | 1E7 |
| $\alpha$ | Cells-eDNA binding rate | 4E-5 |
| $R_j$ | eDNA degradation rate | $0.15 \forall j$ |
| $M_{input,j}$ | eDNA molecules input | $1E4 \forall j$ |
| $I$ | Mean stress intensity | 0.2 |
| $d$ | Mean stress duration | $d_1$={100,300,500,700,900,1100}<br>$d_2$=300 (Figure 4, line 1), $d_2$=1100 (Figure 4, lines 2 and 3) |
| $f$ | Mean stress frequency | $f_1$={1E-4,3E-4,5E-4,7E-4,9E-4,1.1E-3}<br>$f_2$=5E-4 (Figure 4, line 1), $f_2$=1.1E-3 (Figure 4, lines 2 and 3) |

#### Supplementary Table S2: Transformation event probabilities used in the model.

| | Initial genotype $i$ | Acquired allele | Final genotype $j$ | Random draw | Probability of success of the transformation undertaken $S_{(i,j)}$ |
|---|---|---|---|---|---|
| Experimental design 1 (model of gene transfer, figure 2A) | WT | comM::WT or pilT::WT or pho::WT | WT | $P_1$~N(-3.65,0.11) | $10^{P_1}/10^{P_1}=1$ |
| | WT | comM::AbaR or pilT::AbaR or pho::AbaR | ΔcomM *or ΔpilT or Δpho* | $P_2$~N(-4.27,0.066) | $10^{P_2}/10^{P_1}$ |
| | Δ*pho* | pho::WT | WT | | 1 |
| | Δ*pho* | comM::AbaR or pilT::AbaR | Δ*comM or ΔpilT* | | $10^{P_2}/10^{P_1}$ |
| | Δ*comM* | comM::WT | WT | $P_4$~N(-6.166,0.129) | $10^{P_4}/10^{P_1}$ |
| | Δ*comM* | pilT::AbaR or pho::AbaR | Δ*pilT* or Δ*comM* | $P_5$~N(-7.120.102) | $10^{P_5}/10^{P_1}$ |
| | Δ*pilT* | pilT::WT | WT | | 0 |
| | Δ*pilT* | comM::AbaR or pho::AbaR | Δ*pilT* | | 0 |
| Experimental design 2 (model of gene transfer and allelic transfer, figure 2B) | WT | SNP | WT+SNP | $P_3$~N(-5.246,0.086) | $10^{P_3}/10^{P_3}=1$ |
| | Δ*pho* | SNP | Δ*pho*+SNP | | 1 |
| | Δ*comM* | SNP | Δ*comM*+SNP | $P_6$~N(-5.953,0.091) | $10^{P_6}/10^{P_3}$ |
| | Δ*pilT* | SNP | Δ*pilT*+SNP | | 0 |

The probability of success of the transformation undertaken is equal to 1 when (i) the transformation is not inhibited (i.e. when the cell involved in the transformation is of WT or Δ*pho* genotype) and (ii) the cell acquires a short allele (WT or SNP). In other situations, the probability of a successful transformation is lower, either because of the high length of the acquired allele (AbaR acquisition), or because the *comM* or *pilT* gene is no longer functional (Δ*comM* or Δ*pho*). The probability of a successful transformation is then calculated as the ratio of the probabilities drawn at random from the normal distributions (mean and SE) obtained from the log-transformed frequencies of the experiments (Figure 2). The 'reference' probability (an estimate of the probability of transformation without inhibition and corresponding to the acquisition of a short sequence), P1 and P3, differs according to experimental design 1 or 2, respectively.

### Supplementary Table S3: Bacterial strains and plasmids used in this study

| Name/Genotype | Source | Comments |
|---|---|---|
| **strains used for the measure of acquisition of a single nucleotide polymorphism** | | |
| M2 WT | Niu *et al.* 2008 | *A. nosocomialis* M2 Wild-type strain |
| M2 *comM::[sacB aacC4]* | This study | Sucrose sensitive and apramycin resistant |
| M2 *ΔcomM* | This study | Deletion into the *comM* gene |
| M2 *pho::[sacB aacC4]* | This study | Sucrose sensitive and apramycin resistant |
| M2 *Δpho* | This study | Deletion into the *pho* gene |
| M2 *attn7::[lacZ aphA]* | This study | Natural transformation with assembly PCR |
| M2 *ΔcomM attn7::[lacZ aphA]* | This study | Kanamycin resistant |
| M2 *Δpho attn7::[lacZ aphA]* | This study | Kanamycin resistant |
| M2 *pilT::km* | Wilharm et al., 2013 | Not naturally transformable ; Kanamycin resistant |
| M2 *rpoB*(Rif$^R$) | This study | Spontaneous mutation in *rpoB* gene ; Rifampicin resistant |
| M2 *rpoB*(Rif$^R$) *comEC::aacC4* | This study | Rifampicin resistant, apramycin resistant |
| **Strains used for bacterial competitions** | | |
| M2 *hu-sfgfp aacC4* | Godeux et al., 2018 | Fluorescent ; apramycin resistant |
| M2 *comM::[*AbaR4*]* | This study | Natural transformation with genomic DNA from *A. baumannii* 40288 strain |
| M2 *comM::[*AbaR1*]* | This study | Natural transformation with genomic DNA from *A. baumannii* AYE strain |
| AYE WT | Fournier *et al.* 2006 | *A. baumannii* AYE Wild-type strain |
| M2 *comM::[*AbaR4*] hu-sfgfp aacC4* | This study | Fluorescent ; apramycin resistant |
| M2 *comM::[*AbaR1*] hu-sfgfp aacC4* | This study | Fluorescent ; apramycin resistant |
| M2 *ΔcomM hu-sfgfp aacC4* | This study | Fluorescent ; apramycin resistant |
| **Strains used for the measure of acquisition and deletion of the *L* fragment** | | |
| M2 *pho::[*AbaR4*]* | This study | Natural transformation with genomic DNA from *A. baumannii* AB0057 strain |
| M2 *comM::[sacB aacC4] pho::[*AbaR4*]* | This study | Sucrose sensitive and apramycin resistant |
| M2 *ΔcomM pho::[AbaR4]* | This study | Deletion into the *comM* gene |
| M2 *ΔcomM pho::[AbaR4 sacB aacC4]* | This study | Intermediate genotype |
| M2 *ΔcomM pho::[*AbaR4 *Δ'aacC4]* | This study | Intermediate genotype |
| M2 *ΔcomM pho::[*AbaR4 *tniA::(sacB aacC4) Δ'aacC4]* | This study | Intermediate genotype |
| M2 *ΔcomM pho::[*AbaR4 *tniA1(Oc) Δ'aacC4]* | This study | Intermediate genotype |
| M2 *ΔcomM pho::[AbaR4 tniA1(Oc) aacC4::(aphA Lj)]* | This study | Intermediate genotype |
| M2 *rpoB*(Rif$^R$) *ΔcomM pho::[AbaR4 tniA1(Oc) aacC4::(aphA Lj)]* | This study | Recipient strain for deletion ; deletion into the *comM* gene ; Rifampicin resistant |
| M2 *comEC::tetA ΔcomM pho::[*AbaR4 *tniA1(Oc) aacC4::(aphA Lj)]* | This study | Donor strain for acquisition ; not naturally transformable |
| M2 *ΔcomM pho::[*AbaR4 *tniA1(Oc) aacC4 ΔLj]* | This study | Intermediate genotype |
| M2 *rpoB*(Rif$^R$) *ΔcomM pho::[*AbaR4 *tniA1(Oc)* | This study | Recipient strain for acquisition ; deletion into the *comM* |

| | | |
|---|---|---|
| *aacC4 ΔLj]* | | gene ; Rifampicin resistant |
| M2 *comEC::tetA ΔcomM pho::[*AbaR4 *tniA1(Oc) aacC4 ΔLj]* | This study | Donor strain for deletion ; not naturally transformable |
| M2 *pho::[AbaR tniA1(Oc) aacC4::(aphA L)]* | This study | Intermediate genotype |
| M2 *rpoB*(Rif$^R$) *comM$^+$ pho::[*AbaR4 *tniA1(Oc) aacC4::(aphA Lj)]* | This study | Recipient strain for deletion ; rifampicin resistant |
| M2 *comM$^+$ pho::[*AbaR4 *tniA1(Oc) aacC4 ΔLj]* | This study | Intermediate genotype |
| M2 *rpoB*(Rif$^R$) *comM$^+$ pho::[*AbaR4 *tniA1(Oc) aacC4 ΔLj]* | This study | Recipient strain for acquisition ; Rifampicin resistant |
| **Plasmids** | | |
| pASG-4 | Godeux *et al.* 2018 | Not replicative in *Acinetobacter* |
| pMHL-7 | | Contains the [*sacB aacC4*] cassette |
| pMHL-2 | Godeux *et al.* 2018 | Contains the *aacC4* gene |
| pXDC116 | Derivative of pXDC61 (Charpentier *et al.* 2009) | Contains the Kan$^R$ cassette |

#### Supplementary Table S4: Primers used in this study

| Primer name | Sequence 5' to 3' | Template ; annealing site | Use (genetic construct) |
|---|---|---|---|
| mlo-104 | AAGATATCGGTCTCCAAGC | M2 genomic DNA ; *rpoB* gene | *rpoB*(Rif$^R$) |
| mlo-105 | AGTACGGCCTTCGTCAT | M2 genomic DNA ; *rpoB* gene | *rpoB*(Rif$^R$) |
| mlo-97 | ATACCGCCGTAGAATGCC | M2 genomic DNA ; 2-kbp Upstream *comM* gene | *ΔcomM* |
| comM-Rev | TTAAGAGTGATTACCTCGATAAGA | M2 genomic DNA ; 3' of the *comM* gene | *ΔcomM* |
| asg-100 | AATTTCCAGTGCACGACG | M2 genomic DNA ; *comM* gene | *ΔcomM* |
| asg-76 | CGGCGTGCACTAGAAATTCCACGGGTGAAATTAC | M2 genomic DNA ; *comM* gene | *ΔcomM* |
| Apra-For | ATCAAGGCCCGATCCTTGGAGCCCTTG | pMHL-2 (Godeux et al., 2018) ; Begining of *aacC4* gene | *[aacC4 ΔL] ; aacC4::aphA* |
| mlo-29 | TCATGAGCTCAGCCAATCGACTGG | pMHL-2 (Godeux et al., 2018) ; End of Apra$^R$ cassette | *[aacC4 ΔL] ; [sacB aacC4] ;* [AbaR4 *aacC4::(aphA L)]* (L=2, 4-kbp) |
| asg-58 | ATCACCCATCACATATACCTGCCG | pMHL-2 (Godeux et al., 2018) ; Upstream *sacB* gene | *[sacB aacC4]* |
| asg-85 | ACCCTGTATTTCACGTAG | M2 genomic DNA ; 2-kbp Upstream *pho* gene | *pho::[sacB aacC4] ; pho::[*AbaR4 *sacB aacC4] ; Δpho* |
| asg-86 | CGGCAGGTATATGTGATGGGTGATTACGACGGCTCACATATTGGTC | M2 genomic DNA ; *pho* gene | *pho::[sacB aacC4] ; pho::[*AbaR4 *sacB aacC4] ; Δpho* |
| asg-87 | CCAGTCGATTGGCTGAGCTCATGACTAAAGCAAGCGGTG | M2 genomic DNA ; *pho* gene | *pho::[sacB aacC4]* |
| asg-88 | AACCAGCAATAACACC | M2 genomic DNA ; 2-kbp Downstream *pho* gene | *pho::[sacB aacC4] ; pho::[*AbaR4 *sacB aacC4] ; Δpho ; pho::[*AbaR4 *Δ'aacC4]* |
| asg-97 | ATCACCCATCACATATACCTGCCGTTGGGACGCAATTGG | M2 genomic DNA ; *pho* gene | *Δpho* |
| asg-108 | TTCAGTTGTCTCAAAGGACGC | AbaR4 island | *pho::[*AbaR4 *sacB aacC4] ; pho::[*AbaR4 *Δ'aacC4] ; [*AbaR4 *aacC4::(aphA L)]* (L=2, 10-kbp) |
| asg-137 | CGGCAGGTATATGTGATGGGTGATGTCAGTTCCACAAATAAATCAGAGT | M2 genomic DNA ; Begining of 3'half *pho* gene | *pho::[*AbaR4 *sacB aacC4]* |
| asg-77b | GTCAGTTCCACAAATAAATCAGAGT | M2 genomic DNA ; Begining of 3'half *pho* gene | *pho::[*AbaR4 *Δ'aacC4]* |
| asg-138 | ACTCTGATTTATTTGTGGTGACTCATGCCCTCGTGGTCAG | pMHL-2 (Godeux et al., 2018) ; aacC4 gene | *pho::[*AbaR4 *Δ'aacC4]* |
| mlo-74 | TGACAGAGCTGACGCC | M2 genomic DNA ; Begining of *pho* gene | *tniA::[sacB aacC4] ; tniA1(Oc)* |
| asg-159 | GACCACATTGGGTTATAGAATGATACGACTA | AbaR4 island ; *tniA* gene | *tniA1(Oc)* |

| | | | |
|---|---|---|---|
| asg-160 | TAGTCGTATCATTCTATCCAATGTGGTTGTC | AbaR4 island ; *tniA* gene | *tniA1(Oc)* |
| asg-161 | CGGCAGGTATATGTGATGGGTGATTATTGACAGCATTAGATTC | AbaR4 island ; *tniA* gene | *tniA::[sacB aacC4]* |
| asg-162 | CCAGTCGATTGGCTGAGCTCATGATAGGGCGAATGAATATGATGTG | AbaR4 island ; *tniA* gene | *tniA::[sacB aacC4]* |
| R18 | TGAGAGACGCTACTC | AbaR4 island | *tniA::[sacB aacC4] ; tniA1(Oc)* |
| comEC-For | TGGTGTCGCTGTACTAATTACGGT | M2 genomic DNA ; *comEC* gene | *comEC::aacC4* |
| asg-67 | GGGTTGAGATATACTCGCG | M2 genomic DNA ; 2-kb upstream *comEC* gene | *comEC::tetA* |
| comEC-Rev | CTTCAGTTCACGCATCAAGCTTGT | M2 genomic DNA ; *comEC* gene | *comEC::aacC4 ; comEC::tetA* |
| asg-168b | GGAAGTATCAGAATTGGTTAATCAATGATATGTTGCTCAG | M2 genomic DNA ; *comEC* gene | *comEC::tetA* |
| asg-169 | CATATATAATAATTAAGTTTATTTTTCAAAAAGGCTCAATATACTG | M2 genomic DNA ; *comEC* gene | *comEC::tetA* |
| asg-148 | GACTTGAGGCGCAGCGTCGTCCAGACCTGACC | pMHL-2 ; *aacC4* gene | *aacC4::aphA* |
| kan-F | CTGCGCCTCAAGTGTTC | pXDC116 ; $Kan^R$ cassette | *aacC4::aphA* |
| kan-R | TGCCTCGTGAAGAAGG | pXDC116 ; $Kan^R$ cassette | *aacC4::aphA* |
| R3 | CAAGGGCTCCAAGGATCGGGCCTTGATTTACCATATGTGC | AbaR4 island | *[*AbaR4 *aacC4::(aphA L)]* (L=2-kbp) |
| F4.2 | CCTTCTTCACGAGGCATTAAGGCAAGATTAGC | AbaR4 island | [AbaR aacC4::(aphA L)] (L=2-kbp) |
| oxa23-Rev | ATTTCTGACCGCATTTCCAT | $bla_{OXA-23}$ gene | *[*AbaR4 *aacC4::(aphA L)]* (L=4-kbp) |
| R5 | CAAGGGCTCCAAGGATCGGGCCTTGATTATTCGAGAATGG | AbaR4 island | *[*AbaR4 *aacC4::(aphA L)]* (L=4-kbp) |
| F6.2 | CCTTCTTCACGAGGCTAGACAGTGATGAG | AbaR4 island | *[*AbaR4 *aacC4::(aphA L)]* (L=4-kbp) |
| F7 | TTTGAGACTGAATAAGG | AbaR4 island | *[*AbaR4 *aacC4::(aphA L)]* (L=6-kbp) |
| R7 | CAAGGGCTCCAAGGATCGGGCCTTGATAGATCATCGCAGTAGAC | AbaR4 island | *[*AbaR4 *aacC4::(aphA L)]* (L=6-kbp) |
| F8.2 | CCTTCTTCACGAGGCAAGTTCATTGTCTACTGCG | AbaR4 island | *[*AbaR4 *aacC4::(aphA L)]* (L=6-kbp) |
| asg-107 | TCTTTCGTATCTGAATTTCCACG | AbaR4 island | *[*AbaR4 *aacC4::(aphA L)]* (L=6-kbp) |
| asg-84 | CATTCCAATAAGTTCGACTTCTG | AbaR4 island | *[AbaR4 aacC4::(aphA L)]* (L=8-kbp) |
| R9 | CAAGGGCTCCAAGGATCGGGCCTTGATTGTTACTTGGGTG | AbaR4 island | *[*AbaR4 *aacC4::(aphA L)]* (L=8-kbp) |

| | | | |
|---|---|---|---|
| F10.2 | CCTTCTTCACGAGGCAAGTACCAGAAGC | AbaR4 island | [AbaR4 *aacC4::(aphA L)]* (L=8-kbp) |
| oxa23-D | CAGTGCTTTTAGTTGTTGTGA | *bla*$_{OXA-23}$ gene | [AbaR4 *aacC4::(aphA L)]* (L=8-kbp) |
| F11 | AAAGCGTATCTTGCTG | AbaR4 island | [AbaR4 *aacC4::(aphA L)]* (L=10-kbp) |
| R11 | CAAGGGCTCCAAGGATCGGGCCTTGATTCGCCATGGCAGT G | AbaR4 island | [AbaR4 *aacC4::(aphA L)]* (L=10-kbp) |
| F12.2 | CCTTCTTCACGAGGCAAACTGCCATGGCG | AbaR4 island | [AbaR4 *aacC4::(aphA L)] (*L=10-kbp) |
| F13 | TTTCTCAGATACAGCC | AbaR4 island | [AbaR4 *aacC4::(aphA L)]* (L=12-kbp) |
| R13 | CAAGGGCTCCAAGGATCGGGCCTTGATAAGCACCGTAATTC TC | AbaR4 island | [AbaR4 *aacC4::(aphA L)]* (L=12-kbp) |
| F14.2 | CCTTCTTCACGAGGCAATGAGAATTACGGTGC | AbaR4 island | [AbaR4 *aacC4::(aphA L)]* (L=12-kbp) |
| asg-79 | AGAAGTCGTTATTGG | AbaR4 island | [AbaR4 *aacC4::(aphA L)]* (L=12-kbp) |
| mlo-111 | CACTAGAAGCGCCAAGTACGA | AbaR4 island | [AbaR4 *aacC4::(aphA L)]* (L=14-kbp) |
| R15 | CAAGGGCTCCAAGGATCGGGCCTTGATATCTGAGAGACGC | AbaR4 island | [AbaR4 *aacC4::(aphA L)]* (L=14-kbp) |
| F16.2 | CCTTCTTCACGAGGCATTTCTCAGATACAGCC | AbaR4 island | [AbaR4 *aacC4::(aphA L)]* (L=14-kbp) |
| R16 | ATGGAAGCACCGTAATTC | AbaR4 island | [AbaR4 *aacC4::(aphA L)]* (L=14-kbp) |
| F17 | TGTCATTTACAGCAATAGAATAGAG | AbaR4 island | [AbaR4 *aacC4::(aphA L)]* (L=16-kbp) |
| R17 | CAAGGGCTCCAAGGATCGGGCCTTGATACAAGCTTTACTGA TGTG | AbaR4 island | [AbaR4 *aacC4::(aphA L)]* (L=16-kbp) |
| F18.2 | CCTTCTTCACGAGGCAATCAGTAAAGCTTGTC | AbaR4 island | [AbaR4 *aacC4::(aphA L)]* (L=16-kbp) |
| R18 | TGAGAGACGCTACTC | AbaR4 island | [AbaR4 *aacC4::(aphA L)]* (L=16-kbp) |

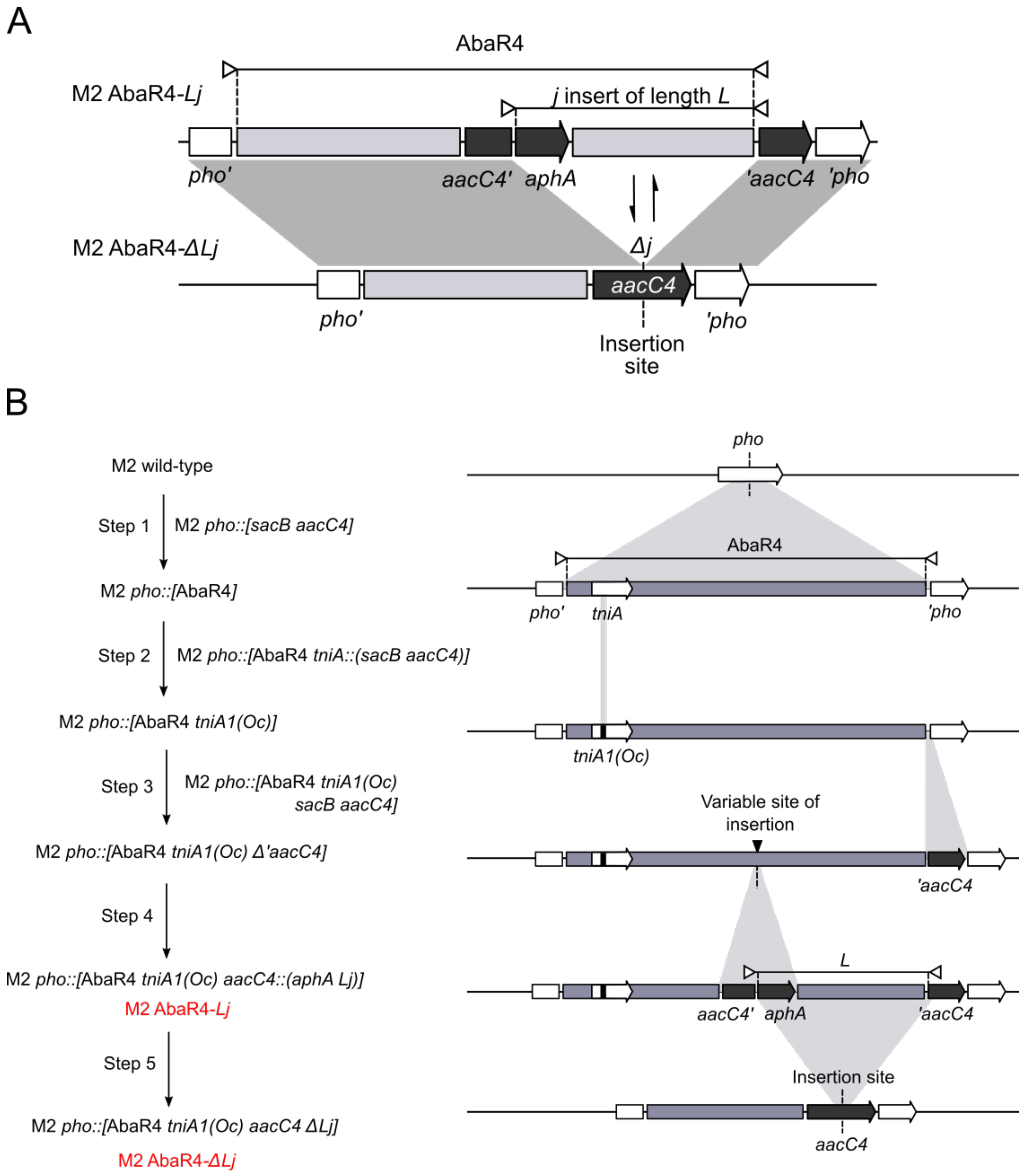

**Supplementary Figure S1. Strategy and mutant construction to quantify events of acquisition or deletion of a heterologous DNA fragment occuring within mixed populations.**

**A. Genotypes of the M2 AbaR4-*Lj*/Δ*Lj* couples used to quantify events of acquisition or deletion.** In the M2 AbaR4-*Lj* genotype, the *j* insert of length *L* is part of an AbaR4 inserted in the *pho* gene, is delimited by the two halves of an *aacC4* apramycin resistance gene (*aacC4'* and *'aacC4*) and contains an *aphA* aminoglycoside resistance gene inserted directly downstream the 3'half of *aacC4*. In the M2 AbaR4-Δ*Lj* genotype, the *j* fragment has been removed to reconstruct the *aacC4* gene. The acquisition of *j* induces the expression of the *aphA* gene and the interruption of the *aacC4* gene leading to a kanamycin resistant/apramycin sensitive phenotype, whereas its deletion induces the reconstruction of an intact *aacC4* gene and the elimination of the *aphA* gene leading to a kanamycin sensitive/apramycin resistant phenotype.

**B. Chromosomal modifications used to generate the genotypes carrying the *j* fragment with a variable length.** The *A. nosocomialis* M2 wild-type strain was modified to insert a large AbaR4 island into the *pho* gene (**step 1**). This was achieved by transforming the M2 *pho::*[*sacB aacC4*] intermediate with purified genomic DNA extracted from the *A. baumannii* AB0057 strain which naturally carries the AbaR4 island inserted into its *pho* gene. To avoid any transposition of the AbaR4 island, a point mutation leading to the generation of a stop codon and a change into the open reading frame was inserted into the coding sequence of the *tniA* gene of the AbaR4, generating the M2 *pho::*[AbaR4 *tniA1*(Oc)] strain (**step 2**). The M2 *pho::*[AbaR4 *tniA1*(Oc) Δ*'aacC4*] strain was then obtained by transforming the M2 *pho::*[AbaR4 *tniA1*(Oc) *sacB aacC4*] intermediate with the 3'half of the *aacC4* gene resulting to its insertion between the end of the AbaR4 and the 3'half of the *pho* gene (**step 3**). This parental strain was then transformed with different genetic constructs composed of the 5'half of the *aacC4* gene and a kanamycin resistance gene (*aphA*) flanked by 2-kbp regions that are homologous to different insertion sites of the AbaR4. It generated 8 different M2 *pho::*[AbaR4 *tniA1*(Oc) *aacC4::*(*aphA Lj*)] genotypes in which an heterologous DNA fragment labelled *j,* had a variable *L* length of 2, 4, 6, 8, 10, 12, 14 and 16-kbp (**step 4**). Finally, each of these genotypes, renamed M2 AbaR-*Lj*, were transformed with a PCR product of the *aacC4* gene to remove the *j* fragment and reconstruct the intact *aacC4* gene (**step 5**). The resulting M2 *pho::*[AbaR4 *tniA1*(Oc) *aacC4* Δ*Lj*)] genotypes were renamed M2 AbaR4-Δ*Lj.* The *ΔcomM* derivatives were obtained by transforming the M2 *ΔcomM* strain with genomic DNA extracted from the different M2 AbaR4-*Lj* and M2 AbaR4-Δ*Lj* strains.

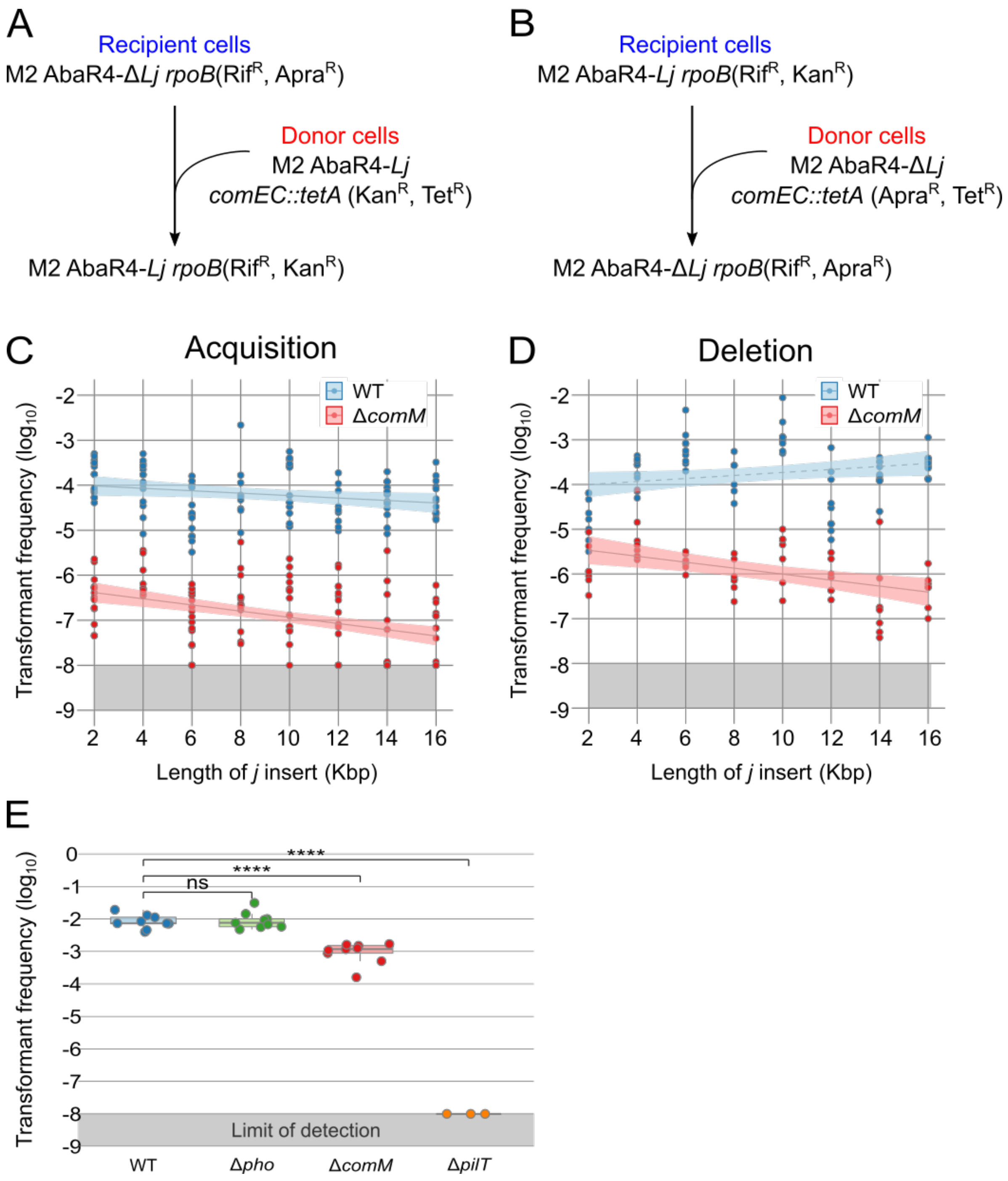

**Supplementary Figure S2. Quantification of rates gene transfer (acquisition and deletion) and rates allelic transfer (SNP) occuring within populations.**

A. and B. Genotypes of derivatives used as recipient or donor to generate the acquisition and the deletion of the *j* fragment. Acquisition events were obtained by naturally transforming M2 AbaR4-Δ*Lj rpoB2*(Rif$^{R}$) recipient cells with non-transformable M2 AbaR4-*Lj comEC::tetA* and deletion events were obtained by naturally transforming M2 AbaR4-*Lj rpoB2*(Rif$^{R}$) recipient cells with non-transformable M2 AbaR4-Δ*Lj comEC::tetA*. Natural transformation frequencies of acquisition (C) or deletion (C) of a heterologous *j* fragment with variable length in mixed culture. Transformant frequencies are represented using a log transformation of the ratio of the number of transformants to the number of total recipient cells. The genetic backgrounds of the *comM* gene are represented for acquisition and for deletion of the *j* fragment (*comM*$^{+}$ in black and Δ*comM* in blue). Detection limit (*F*=-8) is indicated by the light grey area. Experimental events considered as under this limit and were given the detection limit value. A regression linear model is represented for each condition, as a full line when the slope is significantly different from 0 or as a pointed line if not. E. Transformation rates of a single mutation in the *rpoB* gene were measured for the wild-type strain (WT), a mutant with a deletion in the *comM* gene (Δ*comM*), a mutant with a deletion in the *pho* gene (Δ*pho*) and a non-transforming mutant (*pilT::aphA*). Transformation assays were performed using purified genomic DNA extracted from a M2 *rpoB2* ($Rif^R$) *comEC::aacC4* donor. Transformant frequencies represent the ratio between the number of transformants and the total number of recipient cells. The limit of detection for transformants frequencies is indicated by the grey area ($F < 10^{-8}$). Data are shown as experimental replicates from at least 4 independent experiments. Statistics: Mann-Whitney non-parametric test, ****$P$<0.0001.

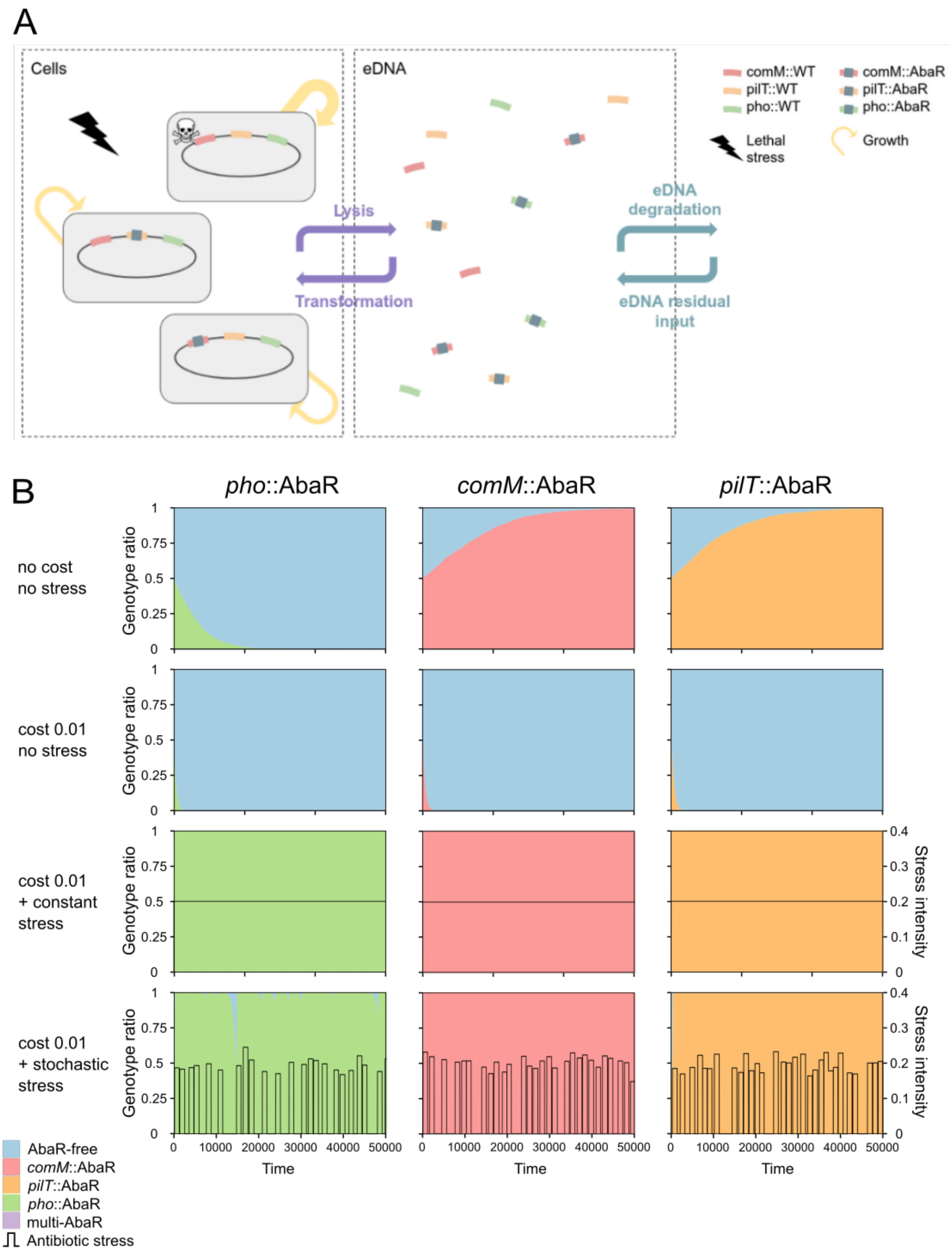

**Supplementary Figure S3. A mathematical model of *A. baumannii* population involving HGT by natural transformation.**

A. Schematic representation of the stochastic computational model for modeling AbaR dynamics within a bacterial population. The model comprises two compartments, one composed of bacterial cells, the other of extracellular DNA (eDNA). The bacterial population grows according to a logistic model. The bacterial cells have three AbaR insertion site in their chromosome (*comM*, *pilT*, *pho*) at which two types of alleles from the eDNA compartment can be integrated by transformation and replace their current DNA: wild-type allele and AbaR carrying resistance (*comM::AbaR*, *pilT::AbaR*, *pho::AbaR*). The integration of a WT allele is costless for cells, whereas the integration of AbaR causes a decrease of cell replication. Bacterial populations are faced with stochastic stresses of random duration, frequency and intensity. In the absence of stress, cells are lysed at a basal rate. Under stress exposure, the lysis rate of WT cells increases but remains unchanged for cells with an AbaR carrying resistance. Each lysed cell releases its DNA and fuels the extracellular compartment with eDNA. AbaR and WT alleles are constantly added to the extracellular environment at a marginal rate simulating residual arrival from neighboring populations. The WT alleles and AbaR are degraded at a constant rate in the extracellular compartment.

B. Examples of simuled dynamics of bacterial genotypes carrying AbaR in competition with AbaR-free cells (wild-type) represented as Mueller plots. Simulations were initialized with equal number of AbaR-free cells and AbaR-carrying cells (*comM*::AbaR, *pilT*::AbaR or *pho*::AbaR). First row, when AbaRs carry no cost and confer no antibiotic resistance. Second row, when AbaRs carry a cost on fitness (0.01) and confer resistance to a stress, but no stress is applied. Third and forth rows, same as in second row, but a stress is applied either continuously or stochastically.

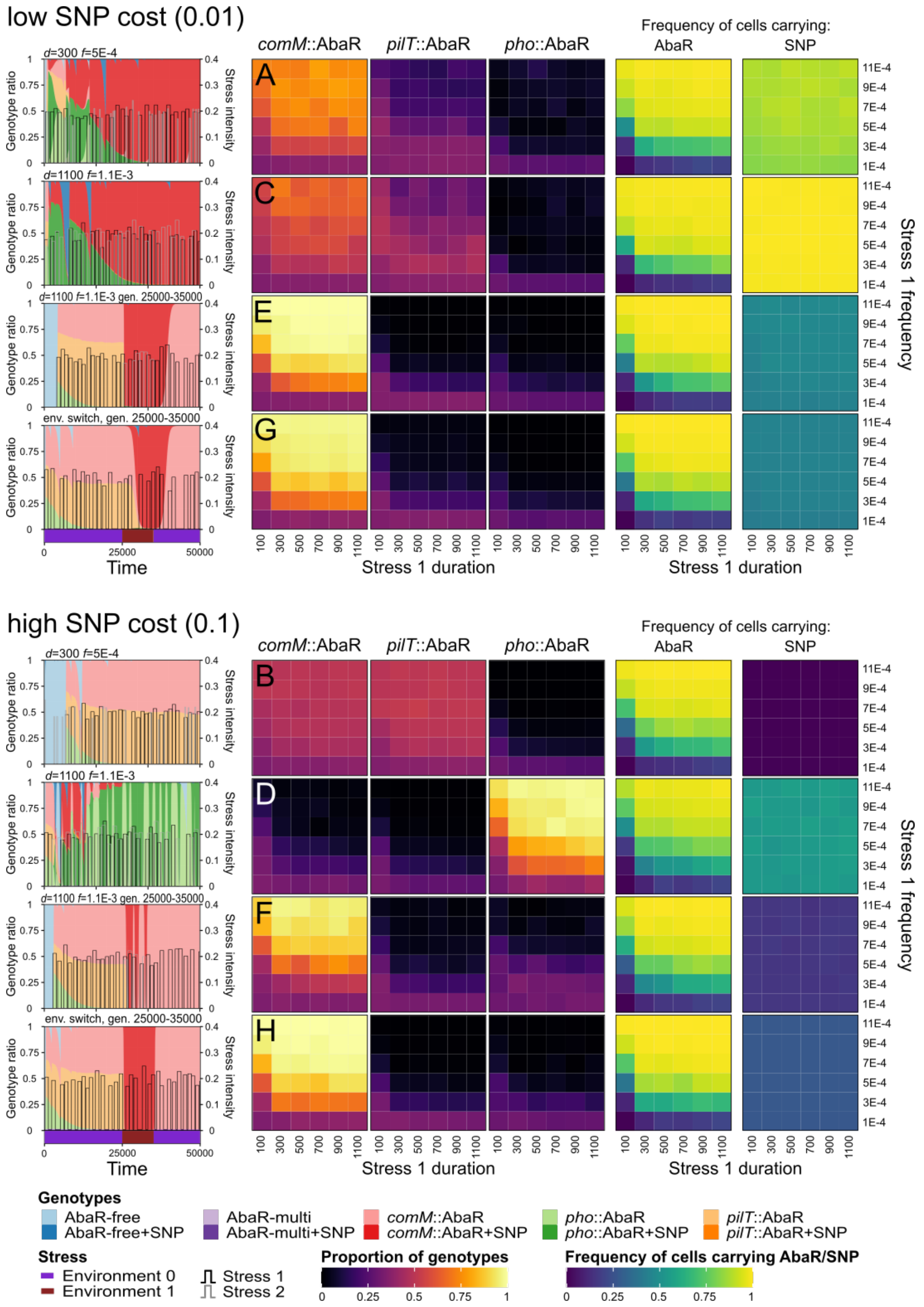

**Supplementary Figure S4. Temporal dynamics and frequencies of bacterial genotypes carrying AbaR.** Temporal dynamics of bacterial genotypes carrying AbaR in environments exposed to a stochastic stress 1 (to which resistance is conferred by AbaR) and another stress 2 to which resistance is conferred by a SNP acquired by allelic transfer. Dynamics were simulated when the SNP is associated to a cost identical to that of AbaR (low cost, 0.01; top panel) and to a much higher cost (high cost 0.1; bottom panel). In each panel, the first column show examples of dynamics (Mueller plot) of the AbaR-carrying genotypes and carrying (+SNP) or not the SNP conferring resistance to stress 2 (or adaptation to environment 1). Stress 1 and stress 2 are displayed as black and gray lines, respectively. Purple and brown boxes below the bottom Mueller plot represent environment 0 and 1, respectively. From top to bottom rows stress 2 are rare and short (mean peak duration *d*=300, mean peak frequency *f*=5E-4), frequent and long (mean peak duration *d*=1100, mean peak frequency *f*=1.1E-3), limited to a short interval (25000-35000) and lastly, when stress 2 is a non-lethal change in environment randomly occurring between 25000 and 35000 time units. Mean frequencies of each AbaR genotype between times 30000 and 50000 were plotted as a function of stress 1 duration and stress 1 frequency (averaged from 100 simulations each). Frequencies of cells carrying AbaR (irrespectively of their insertion site) and frequencies of cells carrying SNP were also plotted as a function of stress 1 duration and stress 1 frequency.
